# Two SID-1-dependent genes sensitive to heritable epigenetic changes can also impact reproduction

**DOI:** 10.64898/2026.06.08.730950

**Authors:** Aishwarya Sathya, Nathan M. Shugarts Devanapally, Andrew L. Yi, Antony M. Jose

**Affiliations:** Department of Cell Biology and Molecular Genetics, University of Maryland, College Park, MD 20742

**Author notes:** Corresponding author: Antony M. Jose **Email:**. **Author Contributions:** AS, NSD, ALY, and AMJ designed and performed research. AS and AMJ wrote the paper with comments from all authors.

**Keywords:** RNAi, transgenerational epigenetic inheritance, germline, sperm

## Abstract

Import of double-stranded RNA (dsRNA) into the germ line can have consequences that last for many generations. However, the role of such transgenerational regulation by extracellular dsRNA is unclear. In the nematode *C. elegans*, entry of dsRNA into the cytosol requires the transmembrane protein SID-1 and loss of SID-1 for a few generations causes changes in gene expression that can persist for hundreds of generations. Here we report an expanded number of such SID-1-dependent genes (SDGs) and analyze two germline-expressed SDGs: *sdg-1* and *sdg-2*. Deleting *sdg-1* reduces brood size in some lineages. An endogenous SDG-1::mCherry fusion protein shows conditional enrichment within nuclei, colocalization with perinuclear germ granules, and colocalization with microtubules. Although animals with SDG-1::mCherry have a normal brood size, they have fewer early progeny with some animals showing defective germline morphology. Deleting the *sdg-1* open reading frame eliminates defects in most but not all the animals that express mCherry in a now *sdg-1(−)* background, suggesting transgenerational consequences of SDG-1::mCherry that persist in some siblings lacking *sdg-1*. Deleting *sdg-2* also reduces brood size in some lineages. An endogenous SDG-2::mCherry fusion protein is constitutively detectable in the cytoplasm and nucleus. The sequence and predicted structure of SDG-2 suggest that it can interact with the Gli-type transcription factor TRA-1, which regulates spermatogenesis. Together, these results suggest that changes in SDG-1 or SDG-2 can impact reproduction. Therefore, the import of extracellular dsRNA or other SID-1 function(s) that regulate SDGs could have evolved to modulate the lingering impacts of ancestral epigenetic changes.

## Introduction

Inheritance of form and function from parent to progeny requires copying DNA sequences and recreating gene expression patterns. Patterns of gene expression can be altered without changes in DNA sequence from one generation to the next when a gene is susceptible to epigenetic changes that are heritable (reviewed in (Madhani 2025; Webster and Phillips 2025)). Such heritable epigenetic changes could impact the development or physiology of organisms across generations. Heritable epigenetic effects mediated by changes in gene expression require that changes in RNAs, proteins, and/or metabolites be integrated into regulatory architectures with positive feedback loops that can sustain changes for many generations (Jose 2020a; 2020b; 2024). Therefore, any process that includes a positive feedback loop could have evolved to accommodate the quantitative and/or qualitative changes needed for observing heritable effects. Analysis of genes that show heritable changes in expression is necessary for defining the processes, if any, that are heritable yet responsive to ancestral changes.

A variety of processes with positive feedback loops have been implicated in sustaining heritable epigenetic changes. In the ciliate *Paramecium aurelia*, changing the handedness of a row of cilia experimentally can result in the perpetuation of the new handedness for more than 300 generations (Beisson and Sonneborn 1965), presumably because each row is made by using the previous row as a template (‘cortical inheritance’ discussed in (Jose 2018)). This process conceivably does not require any changes in gene expression. In the yeast *Cryptococcus neoformans*, the analysis of DNA methylation patterns led to the deduction that ancestral DNA methylation patterns have persisted for millions of years in the absence of the enzyme that catalyzes new methylation patterns *de novo* (Catania et al. 2020). Such changes in methylation patterns likely impact the expression of numerous genes through indirect mechanisms that are difficult to disentangle. While many cases of heritable epigenetic changes have now been documented (some reviewed in (Jose 2020a; Fitz-James and Cavalli 2022; Ishikawa and Schumacher 2025)), underlying mechanisms are largely unknown. The ability to induce heritable epigenetic changes at one or a few genes by engaging RNA-mediated regulation can provide opportunities for rapid elucidation of epigenetic mechanisms (Chey and Jose 2022). Analysis in the nematode *Caenorhabditis elegans* has revealed two such mechanisms that can operate within the germ line and use overlapping machinery (Ketting and Cochella 2020) with the potential for causing transgenerational effects: piwi-interacting RNA (piRNA)-mediated transcript surveillance (Frolows and Ashe 2021) and double-stranded RNA (dsRNA)-mediated gene silencing (Quarato et al. 2022). Downstream of piRNAs and dsRNAs, the required positive feedback loop for maintaining a specific level of gene expression relies on the production of antisense small RNAs using cleaved and stabilized mRNA fragments of the target gene in every generation. While piRNAs are thought to be strictly expressed within the germ line, dsRNAs can be made from many tissues and, through intercellular transport, can cause silencing of matching genes in other tissues (Jose et al. 2009; Ravikumar et al. 2019), including the germ line (Devanapally et al. 2015).

The entry of dsRNA into cells requires the dsRNA-binding transmembrane protein SID-1 (Winston et al. 2002; Feinberg and Hunter 2003; Marré et al. 2016; Wang and Hunter 2017; Shugarts Devanapally et al. 2025) and the interaction between SID-1 and dsRNA is supported by cryogenic electron microscopy (Cryo-EM) structures (Wang et al. 2024; Zhang et al. 2024) with similar structures in the absence of dsRNA for mammalian homologs (Qian et al. 2023; Sun et al. 2023b; Hirano et al. 2024; Liu et al. 2024a; Navratna et al. 2024; Zhang et al. 2024; Zheng et al. 2024). While SID-1-dependent entry of dsRNA into the germ line (Devanapally et al. 2015) and other tissues (Jose et al. 2009; Ravikumar et al. 2019) can be seen when dsRNA is artificially expressed in neurons, the endogenous dsRNAs that have evolved to depend on SID-1 for import into cells and subsequent gene regulation are unknown. Yet, SID-1-dependent changes in mRNA levels have been observed for some genes using genome-edited strains with well-matched genetic backgrounds (Shugarts Devanapally et al. 2025). Of these SID-1-dependent genes (SDGs), two were reported as showing reproducible changes: *sdg-1* and *sdg-2* (Shugarts Devanapally et al. 2025). This discovery raises two key questions: (1) what processes are regulated by SDGs? and (2) how are the heritable changes in SDGs induced and sustained? Analysis of SDGs could provide clues to the processes that have evolved to use extracellular dsRNA-mediated regulation if the SID-1 dependence of SDGs is driven by failure to import endogenous extracellular dsRNAs. Parts of the machinery used for the initiation and maintenance of epigenetic changes caused by experimentally added extracellular dsRNA are being identified (Schreier et al. 2025; Woodhouse et al. 2025). However, a single model that applies for all genes is untenable because controlled exposure to a pulse of dsRNA has revealed differences in the heritable changes induced by such extracellular dsRNA among different genes targeted by the same dsRNA (Devanapally et al. 2021). Notably, a single-copy transgene that is susceptible to transgenerational silencing by extracellular dsRNA is also susceptible to piRNA-mediated silencing induced by mating (Devanapally et al. 2021) and the initiation of such mating-induced silencing is enhanced upon loss of SID-1 (Shugarts Devanapally et al. 2025). Conversely, the *sid-1* mRNA is targeted by piRNAs and disruption of germ granules required for heritable RNA silencing results in silencing of *sid-1* (Dodson and Kennedy 2019; Ouyang et al. 2019). This interaction between dsRNA-mediated and piRNA-mediated gene regulation supports the hypothesis that changes in SID-1 activity perturb piRNA-mediated regulation leading to the cases where SDGs show transgenerational changes in gene expression.

Here we identify additional SDGs, including some genes that show heritable epigenetic changes. By analyzing two such genes, *sdg-1* and *sdg-2*, which are both expressed in the germ line, we reveal the distinct ways that they can impact reproduction. These findings provide insights into the organismal contexts where SID-1-dependent regulation could have evolved in *C. elegans*.

## Materials and Methods

### Strains and oligonucleotides

All strains were cultured on Nematode Growth Medium (NGM) plates seeded with 100 μl of OP50 *E. coli* and maintained at 20°C, intermittently at 15ºC, or at 25ºC to examine impact of higher temperature. AMJ1159, AMJ1217, AMJ1324, AMJ1372, AMJ1577, AMJ1615, AMJ1170, and AMJ1576 created using Cas9-mediated genome editing in (Shugarts Devanapally et al. 2025) were used here. WM49, MDX44, and BB239 were obtained from CGC. All DNA and RNA, except probes for smFISH, were obtained from Integrated DNA Technologies. See Supplementary Table 1 (strains) and Supplementary Table 2 (oligonucleotides).

### Genome editing and transgenesis

For Cas9-mediated genome editing (Dokshin et al. 2018), a general protocol of crRNA (0.056 µg/µl) and tracrRNA (0.1 µg/µl) incubation with the Cas9 enzyme (1 µg/µl) for 10 min. at 37ºC, followed by injection along with homology repair template (200 ng/µl) and pRF4 (40 ng/µl) as a coinjection marker in a 10 µl total mix was used with the specific oligonucleotides and reagents as described below. All plasmids were purified from bacterial culture using QIAprep Spin Miniprep Kit (Qiagen) and all PCR products were generated with Phusion® High-Fidelity DNA Polymerase (New England BioLabs) and purified using NucleoSpin Gel and PCR Clean-up Kit (Macherey-Nagel).

#### To delete *sdg-2* coding sequence

Two crRNAs targeting near the start (P63) and the end (P64) of *sdg-2* coding sequence, tracrRNA, Cas9, a *sdg-2 (deletion)* homology repair template (P65) and pRF4 were injected into N2 animals. After isolating F1 progeny, genotyping for *sdg-2 (deletion)* was performed using the primers P56 and P57 to identify heterozygotes and eventual homozygotes to establish AMJ1347 and two other strains. The *sdg-2* deletion was verified using Sanger sequencing of DNA from each of the three strains.

#### To tag *sdg-2* with *mCherryΔpi*

The crRNA targeting near the *sdg-2* stop codon (P66), tracrRNA, Cas9, an *mCherryΔpi* homology repair template amplified from a plasmid used for MosSCI (Devanapally et al. 2021) with ~30 bp homology with primers P67 and P68, and pRF4 were injected into N2 animals and subsequent screening were performed as described above. Genotyping for *sdg-2::mCherry* was performed using triplex PCR with primers P58, P59 and P60 and the insertion was followed across generations to ultimately isolate AMJ1371 and verify insertion as above.

#### To introduce a premature stop codon into *sid-1*

The crRNA targeting the region near codon encoding Q154 in SID-1 (P77), tracrRNA, Cas9, a homology repair template with the mutation (P78) and pRF4 were injected into AMJ1371 and subsequent screening were performed as above. Genotyping for *sid-1(nonsense)* was performed using PCR with primers P61 and P62, followed by restriction digestion with HpyCH4V to ultimately isolate AMJ1388 and verify the mutation as above.

#### To revert the premature stop codon (Q154Stop) in *sid-1*

The crRNA targeting the region near the premature stop codon (P79), tracrRNA, Cas9, a homology repair template with the reversion (P80) and pRF4 were injected into AMJ1388 and subsequent screening was performed as above. Genotyping for *sid-1(reversion)* was performed using PCR with primers P61 and P62, followed by restriction digestion with HpyCH4V to ultimately isolate AMJ1411 and verify the mutation as above.

#### To express *sdg-2p::mCherryΔpi* from a single-copy transgene

Animals were transformed with plasmids and/or PCR products using microinjection (Mello et al. 1991) to generate extrachromosomal arrays or single-copy transgenes as part of the procedures needed for Mos1-mediated Single Copy Insertion (MosSCI; (Frokjaer-Jensen et al. 2012)). pALY01 (*sdg-2p::mCherryΔpi::sdg-2 3’UTR* with *ttTi5605* homology arms and *Cbr-unc-119(+)*) was generated by amplifying the vector backbone of pCFJ151 (Frokjaer-Jensen et al. 2008) with primers P69 and P70, the *sdg-2* promoter sequence from N2 gDNA with primers P71 and P72, *mCherryΔpi* from a plasmid used for MosSCI (Devanapally et al. 2021) with primers P73 and P74, and the *sdg-2 3’UTR* with primers P75 and P76 and assembling them using NEBuilder® 123HiFi DNA Assembly (New England BioLabs). pALY01 (50 ng/μl) and the coinjection markers pCFJ601 (50 ng/μl), pMA122 (10 ng/μl), pGH8 (10 ng/μl), pCFJ90 (2.5 ng/μl), and pCFJ104 (5 ng/μl) (plasmids described in (Frokjaer-Jensen et al. 2008; Frokjaer-Jensen et al. 2012)) were injected into the germ line of adult EG4322 animals. Three different transgenic lines were isolated as described (Frokjaer-Jensen et al. 2012) and the one used here was designated as AMJ1486. Integration of *sdg-2p::mCherryΔpi::sdg-2 3’UTR* in all three isolates was verified by Sanger sequencing.

### Genetic crosses

#### To create *adr-2(uu28)* animals with *sdg-1(jam232)* in the background

BB239 (*adr-1(−)*; *adr-2(−)*) was crossed with AMJ1577 males followed by re-homozygosing of genotypes to create AMJ1860, AMJ1862 and AMJ1863. Genotyping was performed using triplex PCR with the following primers: P44, P45, and P46 for *adr-2*; P50, P51, and P52 for *sdg-1*; and P47, P48 and P49 for *adr-1*.

#### To create *sdg-1p::sdg-1::mCherryΔpi* animals with *cylc-2::mNG::3xFLAG* in the background

AMJ1372 was crossed with MDX44 males followed by re-homozygosing of genotypes to create AMJ2006. Genotyping was performed using triplex PCR with the following primers: P53, P54, and P55 for *sdg-1::mCherryΔpi*; and P81, P82, and P83 for *cylc-2::mNG::3xFLAG*.

#### To create *sdg-2p::sdg-2::mCherryΔpi* animals with *cylc-2::mNG::3xFLAG* in the background

AMJ1371 was crossed with MDX44 males followed by re-homozygosing of genotypes to create AMJ2109. Genotyping was performed using triplex PCR with the following primers: P58, P59, and P60 for *sdg-2::mCherryΔpi*; and P81, P82, and P83 for *cylc-2::mNG::3xFLAG*.

#### Outcrosses

AMJ1372 was crossed with N2 males followed by re-homozygosing of genotypes to create AMJ2084 and AMJ2085. AMJ1612 was crossed with N2 males followed by re-homozygosing of genotypes to create AMJ2077 and AMJ2078.

### RNAseq and Whole-genome sequencing

The previously generated polyA+ RNAseq data ((Shugarts Devanapally et al. 2025); GSE185385) was re-analyzed to identify additional *sid-1*-dependent genes with potentially smaller effect sizes. While previous analyses (see ‘For analysis of newly generated *sid-1(−)* alleles’ section in (Shugarts Devanapally et al. 2025)) used the Caenorhabditis_elegans.WBcel235.101.gtf annotation file followed by Salmon (Patro et al. 2017) for counting reads that map to a gene, the analyses presented here used the Caenorhabditis_elegans.WBcel235.109.gtf file followed by featureCounts (Liao et al. 2014) and DESeq2 (Love et al. 2014). Also, the previous analyses only used Caenorhabditis_elegans.WBcel235.cdna.all.fa.gz to index for Salmon, which does not include small non-coding RNAs (which are in Caenorhabditis_elegans.WBcel235.ncrna.fa.gz).

In this work, the sorted .bam files generated after mapping using hisat2 (Shugarts Devanapally et al. 2025) were analyzed using a custom R script that implements featureCounts followed by DESeq2 (sid-1_lof_dependent_changes_featureCounts_from_bam.R; R v. 4.5.1). Basic information on all *C. elegans* genes were obtained using SimpleMine as on 25 Feb 2022, merged with the SDGs and presented as a table (Supplementary File 1). A proportional Venn diagram was created using https://www.biovenn.nl/ and adjusted in Adobe Illustrator for display.

The comparison of *sid-1(deletion)* (*sid-1(jam113)*) with its wild type also revealed substantial changes in non-coding RNAs, which are included in Supplementary File 1 and can be seen by removing the filter for biotype (protein-coding gene and pseudogene). The RNA from *sid-1(deletion)* (*sid-1(jam113)*) was sequenced along with RNA from wild type (N2) in one batch, and in a separate batch, a wild type (N2), *sid-1(nonsense)*, and *sid-1(reversion)* were sequenced. Comparison of the wild types used in the two batches showed 2280 genes that changed by >2-fold (code included in sid-1_lof_dependent_changes_featureCounts_from_bam.R), revealing a large batch effect and underscoring the importance of batch-matched controls for identifying *sid-1*-dependent genes. The reasons for this large batch effect could be differences in worm growth (stage, contamination, nutrient availability etc.), in RNA preparation, or in subsequent Illumina sequencing. Therefore, additional experiments are needed to evaluate if these non-coding RNAs are truly *sid-1*-dependent and, like the other genes identified by this comparison, potentially also require certain environmental conditions for SID-1-dependent change.

Genomic DNA was prepared from six strains (N2, AMJ1159, AMJ1217, AMJ1324, AMJ1389, and AMJ1446) and Whole Genome Sequencing was performed using Omega Bioservices. Reads were analyzed using custom scripts (fastq_to_sorted_bam_many.sh followed by sorted_bam_from_bowtie2_to_mutated_genes_for_sid1_crispr.sh,). Briefly, reads were mapped to the *C. elegans* genome (ce11) using Bowtie2 (Langmead and Salzberg 2012) and variants were called using SnpEff (Cingolani et al. 2012). Variants listed for each strain (Supplementary File 2) are those predicted to have a ‘high’ or ‘moderate’ impact and detected in the strains subject to genome editing but not in the wild-type background into which the edits were introduced. Scripts are available on GitHub at https://github.com/AntonyJose-Lab/Sathya_et_al_2026 and the fastq files are available on Sequence Read Archive (PRJNA1475530).

### Reverse transcription and quantitative PCR (RT-qPCR)

RT-qPCR experiments were performed and plotted as described earlier (Shugarts Devanapally et al. 2025) with the following sets of primers for the different genes. Gene-specific RT primers [*tbb-2* (P1), *sid-1* (P2), *sdg-1* (P3), *sdg-2* (P4), *sax-2* (P5), *cls-3* (P6), *fbxa-192* (P7), Y102A5C.5 (P8), and Y102A5C.6 (P9)] followed by nested qPCR primers [*tbb-2* (P10 and P11), *sid-1* (P12 and P13), *sdg-1* (P14 and P15), *sdg-2* (P16 and P17), *sax-2* (P18 and P19), *cls-3* (P20 and P21), *fbxa-192* (P22 and P23), Y102A5C.5 (P24 and P25), and Y102A5C.6 (P26 and P27)] were used.

### Western blotting

Total protein from different strains were prepared, run on an SDS-PAGE and blotted as described earlier (Ravikumar et al. 2019) to detect SDG-1::mCherry and α-tubulin, with the following modifications. Mixed stage animals were washed off three to five crowded but not starved 35 mm plates with M9 buffer and immediately frozen at −80°C. Frozen samples were thawed at room temperature and boiled at 95°C for 15 min. in sample buffer (Tris-HCl, pH 6.8: 0.25 M, SDS: 4.0% (w/v), bromophenol blue: 0.9 mM, β-mercaptoethanol: 16% (v/v; freshly added), and glycerol: 30% (v/v)). Boiled samples were centrifuged at 16,000xg for 10-15 min., and the supernatant was collected for western blot. Proteins were separated on a 14% SDS-PAGE and then blotted onto a nitrocellulose membrane (0.2 µm) using TransBlot™ Turbo Transfer System. The blot was probed for mCherry first, stripped by incubating in 0.2% SDS, 0.1 M Tris, pH 6.8, and 1.4% β-mercaptoethanol (v/v) for 1 h at 65ºC), and then probed for Tubulin. The following primary antibodies were used: mouse anti-α-Tubulin (Sigma: T5168; 1:4000 dilution) and rat anti-mCherry (Thermo Fisher Scientific: M11217; 1:1000 dilution). The following corresponding secondary antibodies were used: Rabbit anti-mouse IgG1 HRP (Sigma: SAB3701171, 1:250 dilution) and goat anti-Rat IgG(H+L) HRP (Thermo Fisher Scientific: 31470, 1:5000 dilution). Blots were developed using chemiluminescence detection reagents (Thermo Fisher Scientific: SuperSignal− West Pico PLUS) as per manufacturer’s instructions and imaged using iBright CL1000 imaging system (Invitrogen).

### Single-molecule fluorescence in situ hybridization (smFISH)

Dissection of gonads, followed by smFISH and DAPI staining was performed as described earlier (Devanapally et al. 2021). Custom Stellaris FISH probes were designed against the exons of *sdg-1* sequence using the Stellaris FISH Probe Designer (Biosearch Technologies; https://oligos.biosearchtech.com/), while avoiding exon-exon junctions to allow for the equivalent detection of both spliced and unspliced transcripts. The probe blend to detect *sdg-1* included 16 exon-specific probes (P28-P43) each tagged with Cy5 dye and antisense to *sdg-1* RNA (Supplementary Fig. 2a). Worms were resuspended and incubated for 5 min. at room temperature in ProLong Glass Antifade Mountant (Thermofisher Scientific, P36982) and prepared for imaging by pipetting them on to a 22 x 22 mm square coverslip (#1.5 thickness) followed by mounting on a microscope slide and allowed to cure for 72 hours. All samples within a single experimental set included control strains and were subjected to identical conditions (e.g. incubation times). Precautions for working with RNA (using RNaseZap (ThermoFisher: AM9780), gloves, and mask) were observed throughout the protocol.

### Imaging

#### Widefield microscopy and quantification

Animals were imaged live in 10 μl of 3 mM levamisole on a 2% agarose pad using a Nikon AZ100 microscope and 128 Photometrics Cool SNAP HQ2 camera. A C-HGFI Intensilight Hg Illuminator was used to excite mCherry (filter cube: 530 to 560 nm excitation, 570 dichroic, and 590 to 650 nm emission). Intensity of mCherry was quantified in FIJI (NIH) as described for SDG-1::mCherry earlier (Shugarts Devanapally et al. 2025). Representative images were adjusted in FIJI (NIH) and/or Adobe Photoshop to identical levels for presentation.

#### Confocal microscopy using SoRa spinning disk

Animals expressing fluorescent reporters were imaged live in 8-10 μl of 5 mM levamisole between a cover slip (#1.5 thickness) and a 2% agarose pad. Imaging was done using a CSU-W1 SoRa (Yokogawa) spinning disc with a 60x oil objective lens on an Evident Scientific SpinSR microscope equipped with ORCA-Fusion BT back-thinned Gen-III sCMOS camera. The mCherry fluorescence was excited using a 561 nm laser, reflected toward the sample by a D405/488/561/640 dichroic mirror, and emitted fluorescence was transmitted through the same dichroic mirror and collected using a B617/73 bandpass emission filter. With the same dichroic mirror, mNeonGreen or DAPI fluorescence was excited using 488 nm or 405 nm lasers with emitted fluorescence collected through B525/50 or B447/60 filters, respectively. Images were processed using FIJI (NIH) as above.

#### Confocal microscopy using Stellaris 8

Imaging of animals for smFISH was performed on a Leica Stellaris 8 FALCON laser Scanning Confocal with a 63x/1.4 NA oil objective lens. DAPI fluorescence was excited using a 405 nm laser and emitted fluorescence was collected between 430nm-586nm. Fluorescence from smFISH probes was detected using a white light laser: Atto565 fluorescence was excited at 563nm and emitted fluorescence was collected between 568nm-750nm; and Cy5 fluorescence was excited at 649 nm and emitted fluorescence was collected between 658nm-740nm. Images were processed using FIJI (NIH) as above.

### Counting Sperm

Animals were collected for DAPI staining at ~11 hours post mid-L4 stage, a time preceding the first egg laying event in wild-type animals (Jose et al. 2007). Animals were fixed and stained with DAPI as described for smFISH above. Sperm nuclei were identified by their compact morphology and location around the spermathecal region (Ward and Carrel 1979), counted manually, and recorded using the Cell Counter plugin (FIJI, NIH). The final sperm count for each animal is reported as the sum of counts from each gonad and the number of embryos *in utero*.

### Brood size and feeding RNAi

Single L4-staged hermaphrodite animals were passaged on to individual NGM plates seeded with OP50 and passaged on to a new seeded plates every 24 hours for 8 days. Four days after each parent was passaged, the numbers of animals that were L4-staged or older were counted. Feeding RNAi targeting *bli-1* was performed as described earlier (Raman et al. 2017).

### Exploration of Protein Sequence and Structure

Protein sequences were obtained from UniProt (UniProt 2025) and/or Wormbase (https://www.wormbase.org/#012-34-5) and BLAST (Altschul et al. 1990) searches were performed to identify related sequences. Protein sequence alignments were performed using Clustal Omega (https://www.ebi.ac.uk/jdispatcher/msa/clustalo) and annotated using JalView (Waterhouse et al. 2009). Gene structures were adapted from UCSC genome browser ((Casper et al. 2026); http://genome.ucsc.edu), where repeat sequences are reported using RepeatMasker. Protein structures were obtained from the AlphaFold database (Jumper et al. 2021; Varadi et al. 2022) or predicted using the AlphaFold3 server (Abramson et al. 2024) and annotated using ChimeraX (Pettersen et al. 2021; Meng et al. 2023). Predicted interactions between proteins were identified using AlphaFold 3 by choosing the model with the highest ipTM scores after 5 runs using distinct seeds. Interactions were highlighted using ChimeraX as described earlier (Lalit and Jose 2025). Structural alignments of highly homologous sequences were performed in ChimeraX using the matchmaker command. For more distantly related structures, Vector Alignment Search Tool (VAST; (Gibrat et al. 1996)) and/or Protein Language Model-based search (PLMSearch; (Liu et al. 2024b)) was used. The predicted structure of SDG-1 (AF-G5ECF6-F1-v6) was used for the search against the entire PDB using VAST (https://www.ncbi.nlm.nih.gov/Structure/VAST/vastsearch.html), which relies on secondary structure matches to identify related structures. The top natural proteins that were predicted to be similar based on the similarity score and the alignment length (Supplementary File 3) were overlayed using iCn3d (Wang et al. 2020) and the resultant PDB files were annotated using ChimeraX (Pettersen et al. 2021; Meng et al. 2023). The protein sequence of SDG-1 was also used to identify the closest related protein using PLMSearch (https://dmiip.sjtu.edu.cn/PLMSearch), which relies on a protein language model to look for structural similarity. The top hit of this search was then used to obtain an alignment score using PLMAlign (https://dmiip.sjtu.edu.cn/PLMAlign) (Liu et al. 2025).

### Statistics and data presentation

For single proportions (e.g., *bli-1* feeding RNAi), Wilson’s estimates were used as before (Raman et al. 2017). For brood size data, Student’s t-test or Mann-Whitney U test (a.k.a Wilcoxon-Rank Sum test) was used. Statistical tests were performed using functions in R (v. 4.6.0) when available and plots were generated using R with adjustments using FIJI (NIH) and Adobe Illustrator (v. 29.8.7) for display. Coding suggestions were obtained using Claude or ChatGPT; and all scripts (RT-qPCR_stats.R, sdg1_broodsize_jitter_stats.R, sdg2_broodsize_jitter_stats.R, and sperm_counts.R) are available on GitHub at https://github.com/AntonyJose-Lab/Sathya_et_al_2026.

## Results

### Temporary loss of SID-1 can cause heritable changes in the expression of multiple genes

To sensitively measure changes in RNA levels caused by changes in *sid-1*, we used genome editing to generate animals with the *sid-1* open reading frame (ORF) deleted (*sid-1(deletion)*), with a premature stop codon (*sid-1(nonsense)*), or with subsequent reversion of the stop codon (*sid-1(reversion)*), and performed poly-A+ RNA sequencing on all these strains and the wild-type strain into which these changes were introduced (Shugarts Devanapally et al. 2025). Our previous analyses of these data identified a small number of *sid-1*-dependent genes (SDGs), with only two showing consistent changes in gene expression in the two different *sid-1* mutants – *sdg-1* and *sdg-2* (Shugarts Devanapally et al. 2025). To identify additional *sid-1*-dependent genes that could be perturbed to a lesser degree, we re-analyzed our data using an updated genome annotation file and a lowered threshold for change (Fig. 1a; see Materials and Methods). This approach resulted in the identification of additional protein-coding genes and/or pseudogenes that showed a log_2_[fold-change] > 0.5 with p-adj < 0.5, including all previously identified genes, resulting in the following totals: 12 in *sid-1(nonsense)*, 96 in *sid-1(deletion)*, and 70 in *sid-1(reversion)*, not including *sid-1* (see Supplementary File 1 for gene lists).

**Fig. 1.**
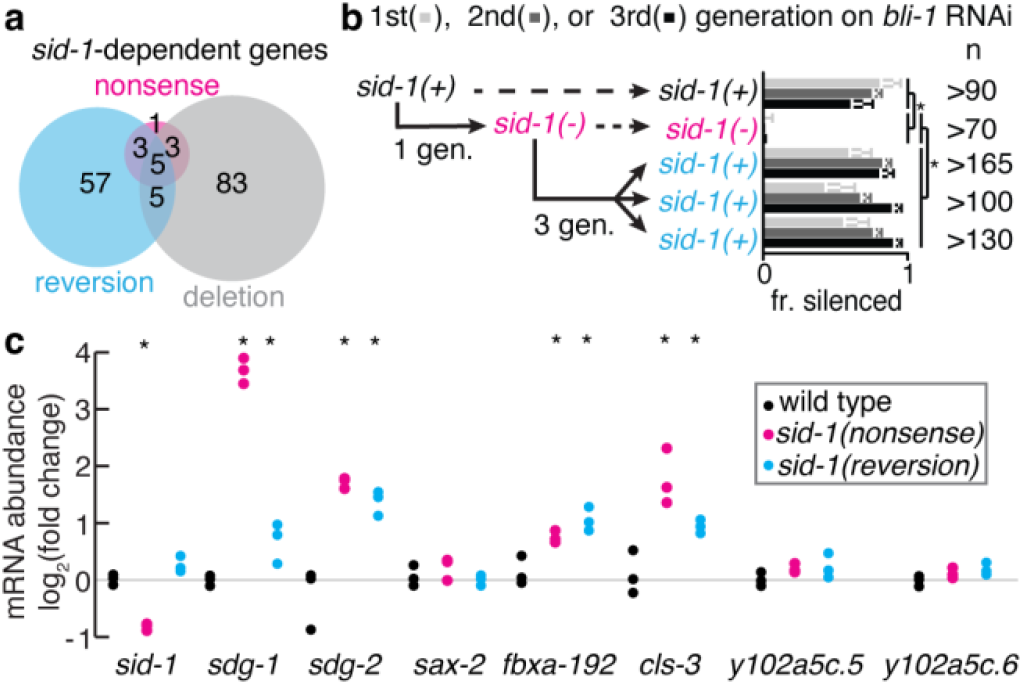
Changes in the expression of some *sid-1*-dependent genes can persist for many generations despite restoration of SID-1 function. a) Numbers of protein-coding genes or pseudogenes that show changes in mRNA levels detected using RNA-seq (abs(log_2_[fold-change]) > 0.5 and p-adj < 0.05) are depicted for *sid-1(nonsense)* [12 total], *sid-1(reversion)* [70 total] and *sid-1(deletion)* [96 total] animals. See Supplementary File 1 for lists of genes. b) Recovery of *bli-1* silencing after reversion of the *sid-1* stop codon using genome editing. Silencing by feeding *bli-1* dsRNA in three successive generations (shades of grey) of wild-type animals (*sid-1(+)*, black), animals with a premature stop codon in *sid-1* introduced through genome editing (*sid-1(−)*, magenta) and three different isolates of animals where the stop codon was reverted to the wild-type sequence using genome editing three generations later (*sid-1(+)*, blue). Error bars indicate 95% CI and asterisks indicate *P < 0*.*05*. c) Abundance of mRNA from *sid-1*-dependent genes as measured using RT-qPCR in wild-type, *sid-1(nonsense)*, and *sid-1(reversion)* animals. Asterisks indicate significant differences (*P < 0*.*05*, Student’s t-test) compared with the corresponding wild type. Values for *sid-1, sdg-1*, and *sdg-2* are replotted here from (Shugarts Devanapally et al. 2025) for comparison.

Both the nonsense mutation in *sid-1* and deletion of *sid-1* result in complete loss of silencing by dsRNA (Shugarts Devanapally et al. 2025). Yet, only 8 genes were identified as having altered mRNA levels in both mutants (Fig. 1a). Furthermore, 5 of these shared genes and 3 other genes remained changed despite reversion of the *sid-1* pre-mature stop codon. Finally, 10 genes were changed in both the deletion mutant and upon reversion of the stop codon. Every overlap is significant (29x to 190x enrichment with *P* < *10*^*-11*^ using a hypergeometric test with Bonferroni correction and considering 20,000 total genes). However, many genes are detected as changed only in the deletion mutant (83 genes) or only in the reversion (57 genes). The presence of many changed genes despite reversion of the nonsense mutation suggests that either these changes reflect heritable epigenetic consequences of a few generations of *sid-1* loss and/or unrelated changes in the background. Our previous detailed analysis of one of these SID-1-dependent genes, *sdg-1*, revealed that changes in its mRNA levels (as measured by RT-qPCR) or changes in its protein levels (as measured using an SDG-1::mCherry endogenous tag) indeed persist after reversion of *sid-1* mutations (Shugarts Devanapally et al. 2025), supporting the possibility of similar susceptibility to heritable epigenetic changes for the genes showing persistent changes here. To test if the observations could be explained by unintended mutations in the background, we performed whole-genome sequencing of multiple strains (Supplemental File 2; *sid-1(nonsense), sid-1(reversion), sid-1(deletion)*, two strains with a *sid-1(nonsense)* in a *sdg-1::mCherryΔpi* background, and the wild-type background into which all these changes were introduced). Fewer than 40 changes that alter protein sequences were detected upon whole-genome sequencing in each of the five strains. Many of these mutations were shared across multiple strains, consistent with pre-existing polymorphisms. The few unique mutations identified in each strain could be the result of differential segregation of pre-existing variation, which is unavoidable. Importantly, none suggest an obvious genetic explanation for the expression changes observed in *sid-1(−)* animals and their persistence despite restoration of wild-type *sid-1* (Fig. 1a). Therefore, we conclude that, as in the case of *sdg-1* (Shugarts Devanapally et al. 2025), some of these *sid-1*-dependent changes in mRNA levels are likely heritable epigenetic changes.

Such heritable epigenetic changes could be caused by persistent losses in dsRNA uptake from the environment despite reversion of the nonsense mutation in *sid-1* or be a separate consequence of the transient loss of SID-1 (e.g., heritable changes triggered upon failure to import endogenous extracellular dsRNA). Therefore, we used silencing of the hypodermal gene *bli-1*, which is a sensitive indicator of RNAi defects (Raman et al. 2017; Knudsen-Palmer et al. 2024), to examine the recovery of dsRNA import from the environment and thus RNAi after restoration of wild-type *sid-1* sequences through genome editing (Fig. 1b). Specifically, we restored wild-type *sid-1* sequence in *sid-1(nonsense)* animals using genome editing and examined *bli-1* RNAi in three successive generations after restoration. Despite a consistent lag observed in the recovery of silencing, RNAi was fully restored within three generations (Fig. 1b), suggesting that the minor changes in SID-1 activity that persist despite reversion are temporary. Therefore, the mRNA changes observed in *sid-1(reversion)* animals are likely the consequence of SID-1-dependent changes that persist independent of SID-1’s ongoing role in the uptake of extracellular dsRNA. We re-tested mRNA levels for a few *sid-1*-dependent genes identified by RNA sequencing using RT-qPCR of newly prepared RNA from populations of wild-type, *sid-1(nonsense)* and *sid-1(reversion)* animals. In addition to the previously reported persistence of changes in *sdg-1* and *sdg-2* levels despite recovery of *sid-1* levels ((Shugarts Devanapally et al. 2025) and replotted in Fig. 1c for comparison), we detected significant and persistent changes in the levels of *fbxa-192* and *cls-3* (Fig. 1c), suggesting that the total list of 157 *sid-1*-dependent genes identified in aggregate by the RNA-seq experiment could include genes in addition to *sdg-1* and *sdg-2* that show heritable epigenetic changes. In all, five protein-coding genes or pseudogenes are changed in *sid-1(nonsense), sid-1(reversion)*, and *sid-1(deletion)* strains as detected using RNA-seq (Supplemental File 1): *sdg-1(sdg-1*.*1 = W09B7*.*2; sdg-1*.*2 = F07B7*.*2) (Shugarts Devanapally et al. 2025), sdg-2(Y102A5C*.*36) (Shugarts Devanapally et al. 2025), sdg-3(R06C1*.*4), sdg-4(Y102A5C*.*5)*, and *cls-3*. Intriguingly, *cls-3* mRNA levels were detected as increased in *sid-1(deletion)* (~5.6-fold) but decreased in *sid-1(nonsense)* (~3.3-fold) and in *sid-1(reversion)* (~3.5-fold) in the initial RNA-seq experiment. However, when the same strains were examined subsequently using RT-qPCR increased levels of *cls-3* mRNA were detected in both *sid-1(nonsense)* and *sid-1(reversion)* animals (~2-3 fold). Similarly, by RNA-seq we detect *fbxa-192* to have increased mRNA levels in both *sid-1(nonsense)* and *sid-1(deletion)* mutants, but not in *sid-1(reversion)* animals; however, subsequent measurement using RT-qPCR, revealed increases in *sid-1(nonsense)* and in *sid-1(reversion)* animals.

One possible explanation for such unstable changes that nevertheless can last for many generations is that loss of SID-1 results in loss of buffering of gene expression across generations as has been proposed (Shugarts Devanapally et al. 2025) and that depending on the individuals that have been selected for passaging, expression can appear stabilized at high or low levels for many generations (see Discussion). Consistently, similar differences in expression were also observed when SDG-1::mCherry was examined in different isolates with *sid-1(nonsense)* mutations after genome editing (Shugarts Devanapally et al. 2025). Also, many genes that change upon mutating SID-1 also change when other regulators of RNA silencing are perturbed (Lalit and Jose 2025), suggesting that their mRNA levels are easily altered. Therefore, we propose that *sid-1* loss changes the mRNA levels of SDGs and that such changes can be stable for many generations despite restoration of *sid-1*, with the direction of change for some SDGs continuing to fluctuate across generations after genetic changes in *sid-1*. Given the challenge in pursuing such genes that show variations in gene expression, here we limit our analysis of the biological roles of SDGs to the two genes that show apparently stable changes across generations when examined using both RNA-seq and RT-qPCR: *sdg-1* and *sdg-2*.

### SDG-1 is a retrotransposon-encoded gene with a minor role in RNA interference

SDG-1 is annotated to be a ~35 kDa protein encoded by a gene located within a retrotransposon and has two domains with faint structural similarity to other proteins. The predicted *sdg-1* gene is within a copy of the Cer9 retrotransposon that is within a ~40 kb genomic region duplicated in the reference wild-type strain N2. Proteins associated with or derived from retrotransposons could be co-opted to play important roles in an organism (e.g., Syncytin in human placenta (Mi et al. 2000)). The predicted SDG-1 protein sequence has paralogs, ZK262.8 and C03A7.2, which share high DNA sequence similarity with each other and are located within copies of Cer8 retrotransposon sequences (red in Fig. 2a; (Shugarts Devanapally et al. 2025)). Additional genes encoded by retrotransposons include CERV-1 (Sun et al. 2023a), within the active retrotransposon Cer1, EGAP798.1 and ZC411.1 in the *sdg-1*-containing copies of Cer9, and ZK228.1 in the ZK262.8-containing copy of Cer8 (Fig. 2a). Notably, the N-terminal and C-terminal halves of ZK228.1 overlap in predicted structure (Supplementary Fig. 1a) and sequence (Supplementary Fig. 1b) with EGAP798.1 and its neighboring gene ZC411.1, respectively, suggesting possible recent separation or mis-annotation of one gene as two neighboring genes. Therefore, to test if the annotated *sdg-1* gene indeed encodes the predicted SDG-1 protein, we performed western blotting using anti-mCherry antibodies on total proteins from a strain with both copies of the endogenous *sdg-1* genes (*F07B7*.*2* and *W09B7*.*2*) tagged with *mCherry* sequences that lack piRNA binding sites (*mCherryΔpi*). A specific band at ~64 kDa that is consistent with the expected molecular weight of SDG-1::mCherry was detected (Fig. 2b), suggesting that the SDG-1 protein is encoded by the annotated *sdg-1* gene and not as a fusion with any neighboring genes.

**Fig. 2.**
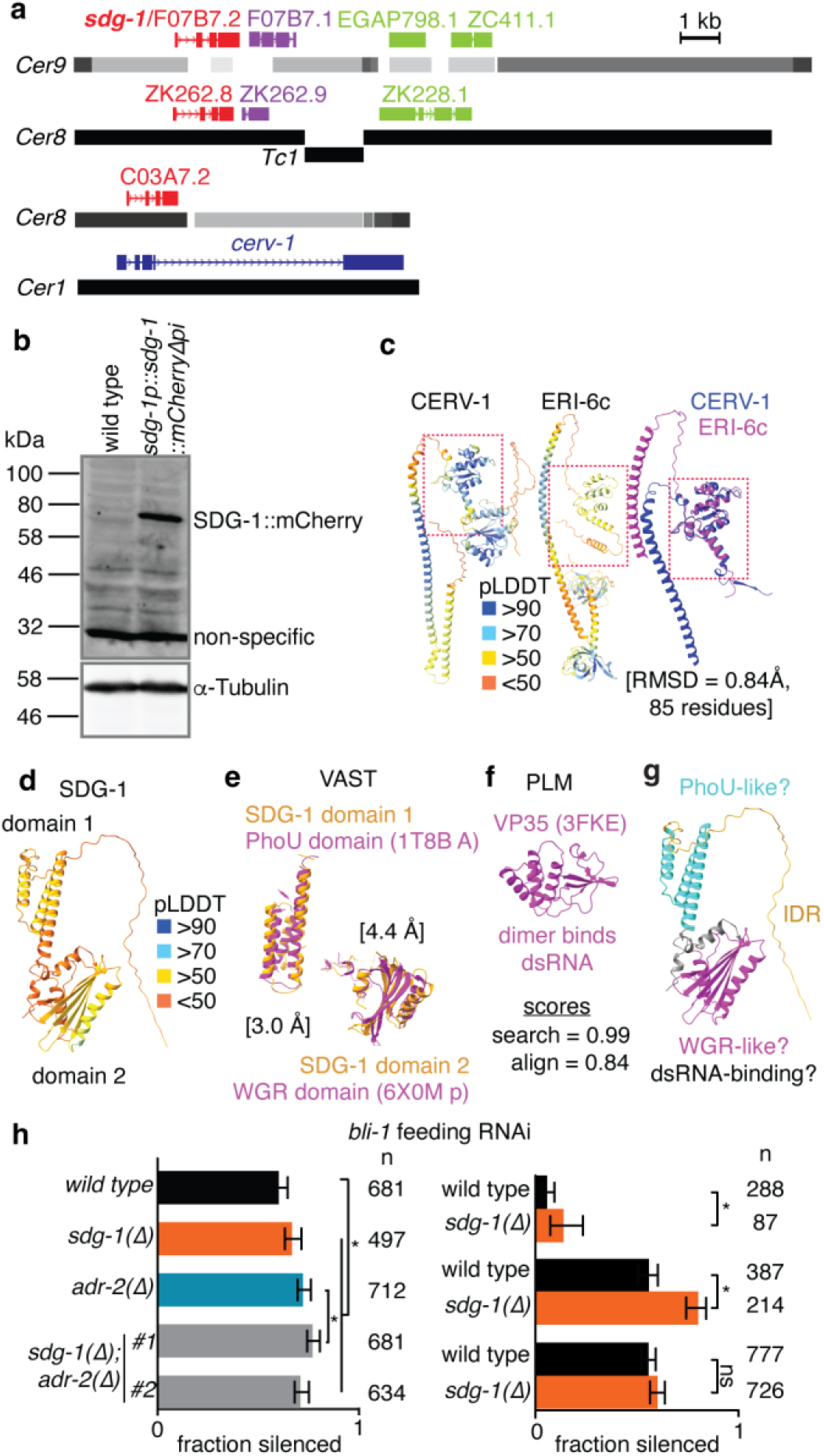
SDG-1 is a retrotransposon-encoded protein with a possible marginal role in RNA interference (RNAi). a) Schematics showing genes encoded within retrotransposons. Similar gene sequences in copies of Cer9, Cer8, and Cer1 are similarly colored, and shades of grey indicate extent of repeat sequences. b) Western blot of total protein extracts from wild-type animals and animals expressing SDG-1::mCherry (*sdg-1p::sdg-1::mCherryΔpi*). A prominent specific band (SDG-1::mCherry, at the expected size of ~64 kDa) and a non-specific band (at ~32 kDa) detected when using anti-mCherry antibodies are indicated. Re-probing for tubulin to control for loading is also indicated (α-Tubulin). c) Predicted structures of CERV-1 and **E**nhanced **R**NA **i**nterference (ERI)-6c along with the overlay of regions with sequence similarity. Left and Middle, Structures of CERV-1 (left) and ERI-6c (middle) as predicted by AlphaFold, colored by confidence scores (pLDDT). Right, Overlay of the predicted structures of CERV-1 and ERI-6c for the regions with sequence similarity (root mean square deviation (RMSD) and aligned number of residues are indicated). Boxes indicate aligned region. See Supplementary Fig. 1c for alignment of CERV-1 and ERI-6c sequences. d) Structure of SDG-1 as predicted by AlphaFold, colored by confidence scores (pLDDT). Note, overall low confidence in predicted structure. e) A PhoU domain, which is found in **pho**sphate regulatory proteins, and a WGR domain, which is rich in Tryptophan (W) Glycine (G) and Arginine (R), and found in poly-[ADP ribose] polymerases, are identified as domains (magenta) similar to the predicted structure of two ordered domains in SDG-1 (orange) as detected using Vector Alignment Search Tool (VAST) similarity search. Numbers indicate RMSD of aligned regions. f) Structure of VP35, which can bind dsRNA as dimers, and is predicted to be close to SDG-1 in the representation used by a Protein Language Model (PLM)-based search with the similarity score (search) and subsequent alignment score (align) indicated. g) Predicted structure of SDG-1 colored by the two predicted domains and an intrinsically disordered region (IDR). h) Impact of *sdg-1* loss on feeding RNAi. Silencing of *bli-1* in wild-type, *sdg-1(Δ), adr-2(Δ)* and two isolates of *sdg-1(Δ); adr-2(Δ)* double mutants are shown (fraction silenced). Right, Trials of *bli-1* feeding RNAi of differing silencing potency for wild-type and *sdg-1(Δ)* animals. Asterisks and error bars are as in Fig. 1b and ns indicates not significant.

Reports on other retrotransposon-encoded genes and the predicted structure of SDG-1 suggest a role for SDG-1 in RNA silencing. The SDG-1 paralog ZK262.8 was identified as causing lethality upon RNAi using a genome-wide RNAi screen in *alg-2(−)* animals (Tops et al. 2006), suggesting a role for ZK262.8 in development. A region of CERV-1 has high structural (Fig. 2c) and sequence (Supplementary Fig. 1c) similarity with the RNA silencing protein Enhancer of RNAi-6c (ERI-6c) (Fig. 2c), suggesting a role for CERV-1 in modulating RNAi. SDG-1 was reported to be enriched near the Z-granule component ZNFX-1 in some animals (Shugarts Devanapally et al. 2025) and to co-immunoprecipitate with the Z-granule surface protein ZSP-1/PID-2 (Placentino et al. 2021), suggestive of interaction with proteins and/or RNAs relevant for RNA silencing. However, the AlphaFold database predicts a structure with low confidence for SDG-1 (Fig. 2d, left; low pLDDT), consistent with the lack of any homologs outside of *C. elegans*. Searching for related structures using the Vector Alignment Search Tool (VAST) identified two domains with some similarity to known protein structures (Fig. 2e) and searching using a protein language model (PLMSearch (Liu et al. 2024b)) suggests that the structure is embedded close to that of the dsRNA-binding protein VP35 from Ebola virus (Fig. 2f). Together, these observations hint at a possible role for SDG-1 in interacting through three distinct domains with other regulators, including potentially dsRNA (Fig. 2g). Our previous deletion of both copies of the *sdg-1* open-reading frame (F07B7.2 and W09B7.2) did not reveal a detectable defect in RNA silencing within the germ line in response to a pulse of *pos-1* dsRNA (Shugarts Devanapally et al. 2025). Nevertheless, since silencing of *bli-1* is a sensitive indicator of RNAi defects (Raman et al. 2017; Knudsen-Palmer et al. 2024), we re-examined animals lacking *sdg-1* in a wild-type background or in a background where RNAi is enhanced upon loss of the dsRNA-editing enzyme ADR-2. We detected a weak but statistically significant enhancement of RNAi upon loss of *sdg-1* in both backgrounds (Fig. 2h, left) and in two of three additional independent comparisons of wild-type and *sdg-1(deletion)* animals with differing potencies of *bli-1* RNAi as evidenced by different extents of silencing in wild-type animals (Fig. 2h, right). Therefore, we conclude that loss of *sdg-1* does not result in a dramatic impact on RNA silencing, although there could be a minor role in silencing by ingested dsRNA.

### SDG-1::mCherry shows dynamic localization during gametogenesis and in early embryos

The SDG-1::mCherry fusion protein previously generated by endogenous tagging is translated from the *sdg-1::mCherryΔpi* mRNA, which is ~16x more abundant in these genome-edited animals than *sdg-1* mRNA in wild-type animals as measured using RT-qPCR (Shugarts Devanapally et al. 2025). Therefore, the relative levels of SDG-1::mCherry protein and whether it impacts SDG-1 function is unclear. As an alternative method of measuring *sdg-1* expression in intact animals, we used smFISH to detect *sdg-1* mRNA in the germ line. However, *sdg-1* transcripts were not detectable above background levels despite detection of control SL1 transcripts and assaying *rde-4(−)* animals (Supplementary Fig. 2), where the level of *sdg-1* mRNA transcripts is expected to be elevated (Welker et al. 2007). Therefore, we proceeded to analyze animals with the SDG-1::mCherry fusion protein, with the caveat that effects seen in animals with high levels of *sdg-1* mRNA, and thus possibly high levels of SDG-1::mCherry fusion protein, could be different from animals with high levels of SDG-1 protein.

Consistent with our earlier findings, SDG-1::mCherry is present in the hermaphrodite germ line and in early embryos (schematic in Fig. 3a, left), where it is excluded from non-dividing nuclei until nuclear envelope breakdown in the −1 oocytes that get fertilized (Shugarts Devanapally et al. 2025). Soon after fertilization, exclusion from nuclei is again detectable in 1-cell embryos before the maternal and paternal pronuclei meet (Supplementary Fig. 3, left). This exclusion is dependent on the presence of the SDG-1 protein because removing the *sdg-1* ORF (Supplementary Fig. 3, middle) or expressing mCherry alone under a different germline promoter (Supplementary Fig. 3, right) results in persistent mCherry enrichment within both parental pronuclei. While fluorescence from SDG-1::mCherry is brightest in the syncytial germ line, we also detected faint fluorescence in the hermaphrodite sperm (Fig. 3a, compare left and middle). Deletion of the *sdg-1* ORF, resulting in animals that express mCherry under the control of *sdg-1 cis*-regulatory sequences, revealed bright fluorescence in sperm (Fig. 3a, right). Marking the spermatocytes and sperm (schematic of spermatogenesis in Supplementary Fig. 4a, top left) with CYLC-2::mNeonGreen (Krauchunas et al. 2020) confirmed that the faint fluorescence is SDG-1::mCherry localized in cell bodies (Supplementary Fig. 4a, top right). Examining spermatogenesis in L4-staged animals revealed that SDG-1::mCherry is abundant in developing spermatocytes, with most of the protein eliminated from sperm along with residual bodies, leaving behind faint fluorescence in spermatids (Supplementary Fig. 4a, bottom). Male germ lines showed a similar progression (Supplementary Fig. 4b). When male spermatids were transferred into hermaphrodites upon mating, SDG-1::mCherry was restricted to the cell bodies and not detected in the pseudopods that form upon activation (Supplementary Fig. 4c). Thus, the SDG-1 protein is detectable in hermaphrodite and male germ lines, with a drastic reduction in developed sperm.

**Fig. 3.**
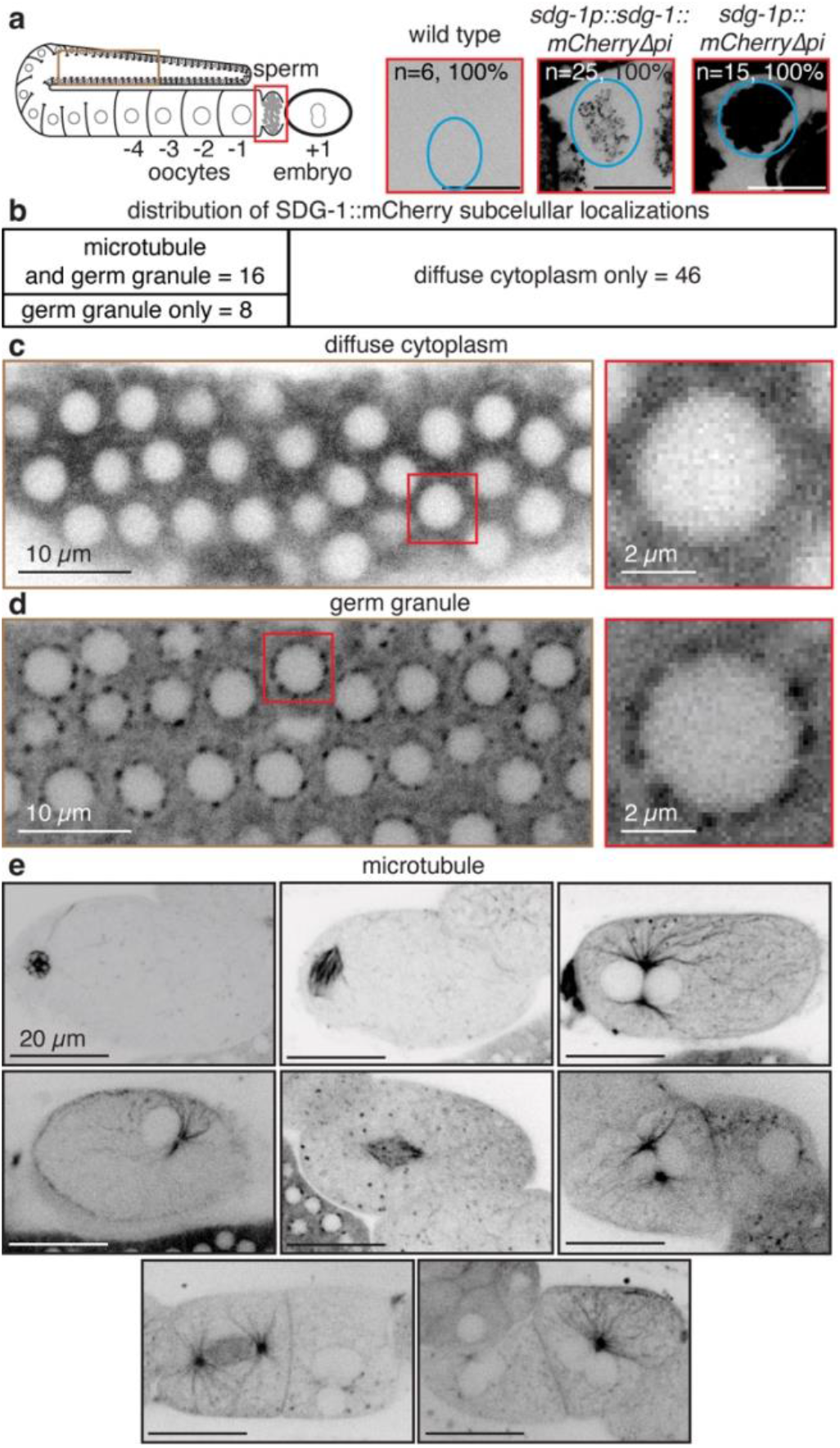
SDG-1::mCherry shows conditional enrichment on germ granules and microtubules. a) Left, Schematic of one arm of the adult germ line where sperm is stored in the spermatheca (red box) and oocytes continually mature (−4 to −1) before fertilization to generate an embryo (+1). Right, Fluorescence (black) in sperm (blue oval) of wild-type animals (left), animals with SDG-1::mCherry (middle) or animals with mCherry expressed under the native *sdg-1* cis-regulatory regions (right). Scale bars, 20 µm. b) Mosaic plot summarizing relative numbers of observations made for the indicated subcellular localizations of SDG-1::mCherry using a mix of imaging conditions. See Supplementary File 4 for details of different imaging conditions. c) Example of diffuse cytoplasmic localization within the germ line (left), with a zoom of one nucleus (right). d) Example of enrichment in perinuclear germ granules within the germ line (left), with a zoom of one nucleus (right). Also imaged using a different microscope in (Shugarts Devanapally et al. 2025). e) Examples of microtubule-like localization in a rough developmental sequence of early embryos from top left to bottom right. Scale bars for each section are as indicated. See Supplementary Fig. 3 for exclusion of SDG-1::mCherry but not mCherry from pronuclei in embryos.

**Fig. 4.**
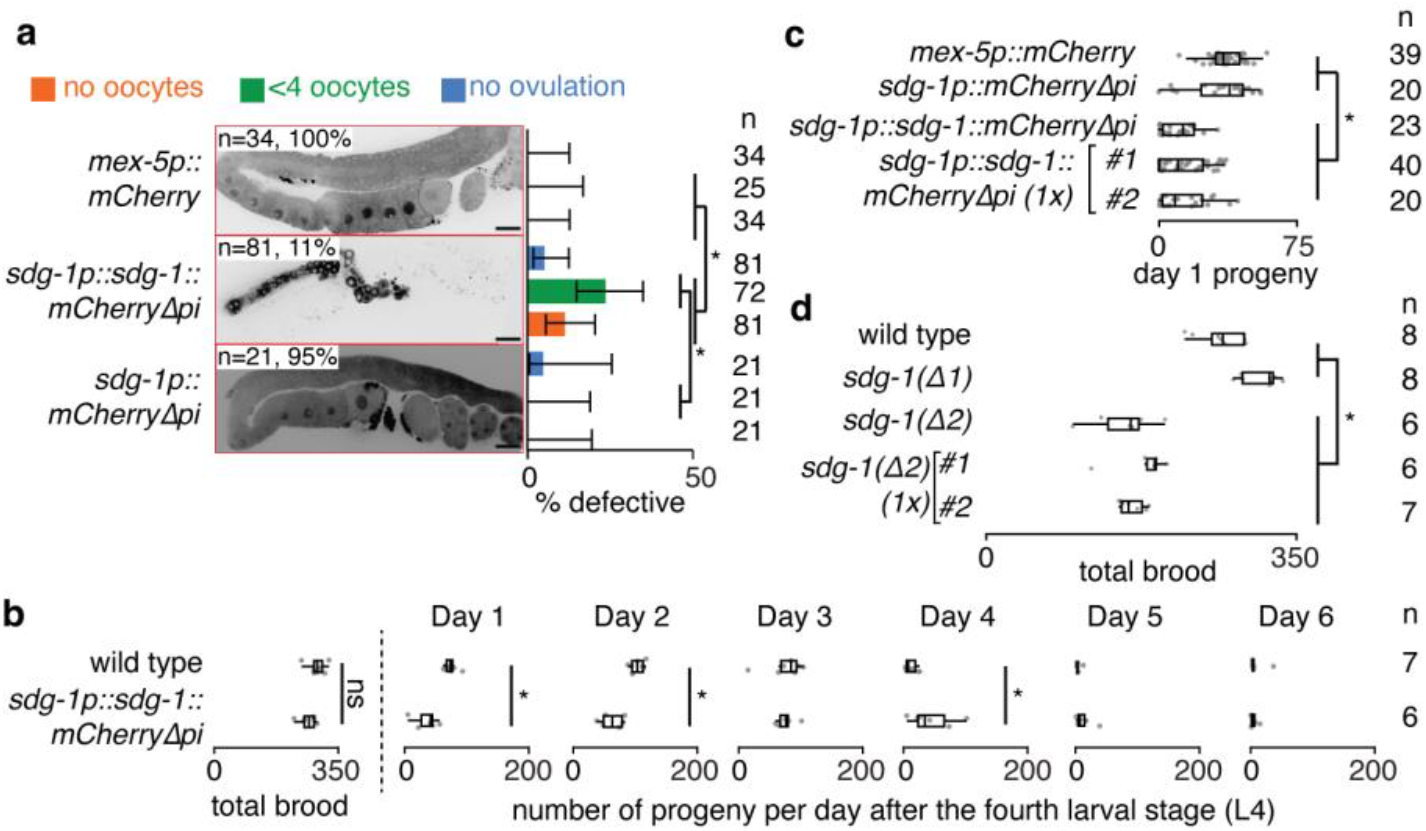
Changes in SDG-1 can impact reproduction. a) Left, Representative gonad arms of young adult animals that express mCherry (*mex-5p::mCherry*) in the germ line, that have SDG-1::mCherry (*sdg-1p::sdg-1::mCherryΔpi*), or that have a subsequent deletion of the *sdg-1* open-reading frame (*sdg-1p:: mCherryΔpi*) are shown. Right, Percentages of defects (categorized as no oocytes, fewer than 4 oocytes, or no ovulation) observed in animals with SDG-1::mCherry are shown for all three strains. Numbers of gonads imaged (n, left) or scorable for each defect (n, right) are indicated. Asterisks and error bars are as in Fig. 1b. Scale bars, 20 µm. See Supplementary Fig. 5 for a gallery of germline defects associated with *sdg-1*. b) Box plot showing total brood and daily cohorts of progeny laid by wild-type animals and animals with SDG-1::mCherry (*sdg-1p::sdg-1::mCherryΔpi*). Significant differences (asterisks; *P < 0*.*05*, Mann-Whitney U test) and numbers of parents analyzed (n) are indicated. c) Box plot showing cohorts of progeny laid in the first day after the fourth larval stage for the indicated strains, including two outcrossed isolates of animals with SDG-1::mCherry. d) Box plot showing total brood size of wild-type animals, two isolates of *sdg-1* deletion (*Δ1* and *Δ2*), and two outcrossed strains (#1 and #2) of the isolate showing a reduced brood size (Δ2). Brood size of *sdg-1(Δ1)* was measured along with the strains in b and plotted here for comparison. Asterisks and n in (c) and (d) are as in (b).

Additionally, we had noted colocalization of SDG-1::mCherry with germ granules marked by ZNFX-1::GFP in ~83% of the animals examined using AiryScan confocal microscopy (Shugarts Devanapally et al. 2025). To examine subcellular localization under gentler imaging conditions without sacrificing resolution, we re-examined animals with SDG-1::mCherry using SoRa spinning disc microscopy. These conditions also revealed the active exclusion of SDG-1::mCherry from non-dividing nuclei, but the enrichment near perinuclear germ granules varied based on imaging conditions (Fig. 3; Supplementary File 4). Specifically, we observed germ granule enrichment most frequently after 30-60 min. incubations in a paralytic or buffer followed by 60-140 min. incubations on a slide before imaging. Strikingly, we also observed localization of SDG-1::mCherry to what appear to be microtubules (Fig. 3), with some cases of multiple spherical structures that exclude SDG-1 within embryos (e.g., last row of embryos in Fig. 3e). In aggregate (Fig. 3b), we observed diffuse cytoplasmic localization with no enrichments in 46 cases (e.g., in Fig. 3c), perinuclear enrichment consistent with germ granules in 24 cases (e.g., in Fig. 3d) and microtubule enrichment in 16 cases (examples in Fig. 3e) in animals where both the germ line and embryos could be imaged in the same frame. Notably, apparent microtubule localization was only observed among animals showing perinuclear enrichments (~66% of such animals). A plausible unifying hypothesis for these observations could be that the subcellular enrichments of SDG-1::mCherry near germ granules and microtubules is in response to imaging conditions. For example, the lack of nutrients for the duration of imaging could have induced enrichment near germ granules, which is a dynamic phase-separated structure with properties that are easily perturbed (Alberti et al. 2019; Uebel and Phillips 2019; Dorner et al. 2026). Additionally, the prolonged pressure on the worm resulting from being sandwiched between an agarose pad and a cover slip could have induced the localization to microtubules. In this light, it is notable that one of the five *sid-1*-dependent genes identified after every *sid-1* change is *cls-3*, which is a homolog of a protein that regulates microtubule dynamics called CLASP (Lawrence et al. 2020). Future experiments are needed to understand the drivers of SDG-1 localization and its biological significance, if any.

### SDG-1 can impact germline development and delay progeny production

When imaging animals with SDG-1::mCherry or with subsequent deletion of the *sdg-1* ORF, we noticed defective germ lines in some animals that appeared either underdeveloped (e.g., Supplementary Fig. 5a) or had abnormal morphology (Supplementary Fig. 5b). Such abnormalities were not observed in animals expressing mCherry alone within the germ line under the control of another promoter (Fig. 4a, compare top and middle). The SDG-1::mCherry fusion protein colocalizes with the Z-granule protein GFP::ZNFX-1 (Shugarts Devanapally et al. 2025), a key protein for heritable gene silencing (Ishidate et al. 2018; Wan et al. 2018), and was identified as a coimmunoprecipitate of another Z-granule protein ZSP-1/PID-2 (Placentino et al. 2021), which is required for maintaining Z-granule structure (Placentino et al. 2021; Wan et al. 2021). Therefore, it is conceivable that perturbation of SDG-1 leads to heritable epigenetic changes that are no longer dependent on the SDG-1 protein. Similarly, some of the defects observed in animals with SDG-1::mCherry could be transgenerational consequences that have become independent of SDG-1. To test this possibility, we compared the germ lines of SDG-1::mCherry animals with those of animals derived from this strain after deleting the *sdg-1* ORF. In such animals that now express mCherry alone under *sdg-1 cis-*regulatory sequences, we no longer observed a reduction in oocytes (Fig. 4a, compare middle with no (orange) or <4 (green) to bottom) 24 hours after the fourth larval stage. Therefore, these defects seen in a total of ~34% of the animals with SDG-1::mCherry can be attributed to the presence of the SDG-1 protein in the assayed animals.

**Fig. 5.**
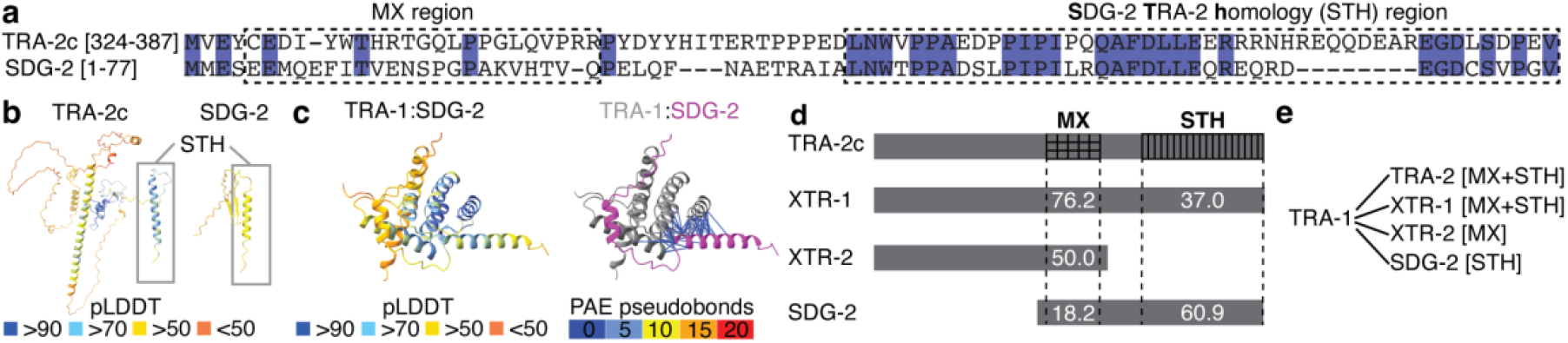
SDG-2 shares homology with the TRA-1-binding region of TRA-2 and is also predicted to interact with TRA-1. a) Alignment of protein sequences of the TRA-2 isoform c (TRA-2c[324-387]) and SDG-2[full length]. Identical residues are highlighted in purple. The previously defined MX region of TRA-2 (left) and the newly defined region of SDG-2 and TRA-2 homology (STH) (right) are boxed. b) Structures of TRA-2c and SDG-2 predicted by Alphafold3 and colored by pLDDT, with the STH region boxed. c) The TRA-1:SDG-2 complex predicted by AlphaFold3 colored by pLDDT (left) or with PAE pseudobonds (right). Terminal disordered regions are not shown. d) Schematic of TRA-1-binding region of TRA-2 and homologous proteins. Identities (%) within the MX and STH domains are indicated. See Supplementary Fig. 7 for sequence alignments. e) Summary of predicted interactions between TRA-1 and TRA-2 or proteins with homology to TRA-2. Interacting domains (MX and/or STH) are in brackets. See Supplementary Fig. 8 for the predicted structures of complexes.

To determine if these defects would lead to a detectable impact on the numbers of progeny, we measured the brood size of these animals along with that of wild-type animals (Fig. 4b). Surprisingly, the total brood was comparable in both strains (Fig. 4b, left). However, the daily cohorts of progeny laid showed significant differences on some days. Animals with SDG-1::mCherry had fewer progeny in the first and second days of egg laying but had more progeny in the fourth day of egg laying, suggesting an apparent delay in the start of egg laying without impact on the total brood. Consistently, this reduction in early progeny was also observed in two outcrossed isolates with SDG-1::mCherry but not in mCherry-expressing control strains or animals lacking the *sdg-1* ORF (Fig. 4c).

When the *sdg-1* ORFs were deleted using genome editing, the three strains obtained with independent deletions had distinct phenotypes: one isolate had no obvious defects (Δ1 in Fig. 4d), one had a reduced brood size that persisted despite outcrossing (compare Δ2 and Δ2 (1x) in Fig. 4d), and a third had a transgenerational Rol defect (potentially related to the use of a plasmid expressing *rol-6* as a co-injection marker) that was seen even in progeny of animals without a detectable Rol defect (Supplementary Fig. 6a), becoming fully penetrant at 25ºC (Supplementary Fig. 6b). While the observed defects could support the idea of multiple SDG-1-dependent heritable epigenetic changes, different background mutations in different genome edited isolates cannot be excluded. Therefore, we conservatively chose the deletion with no apparent defects for examining possible roles of SDG-1 in RNAi (Fig. 2h).

**Fig. 6.**
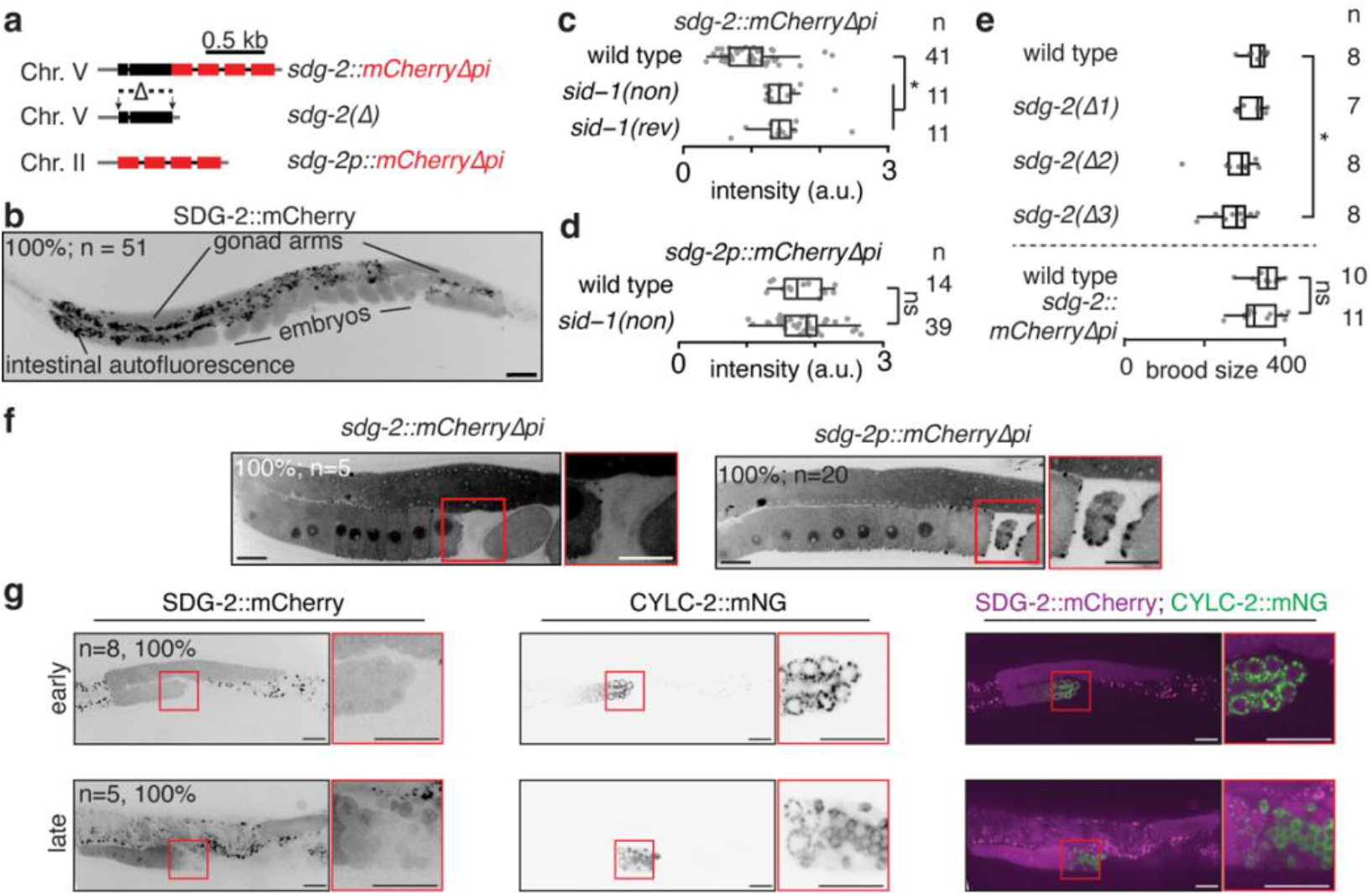
SDG-2 is expressed in the germ line and regulates reproduction. a) Schematic of tools for analyzing the *sdg-2* gene: endogenous tag (*sdg-2::mCherryΔpi*), deletion of the open-reading frame (*sdg-2(Δ)*), and a transcriptional reporter (*sdg-2p::mCherryΔpi*). b) Representative image showing expression of SDG-2::mCherry. Number of animals imaged (n), percentage showing expression pattern, and the punctate intestinal autofluorescence are indicated. Scale bar, 50 µm. c) Box plots showing changes in SDG-2::mCherry fluorescence upon loss of *sid-1* (*sid-1(non)*) or subsequent restoration of wild-type *sid-1* sequence (*sid-1(rev)*). d) Box plots showing SDG-2::mCherry expression in wild-type and *sid-1(non)* animals. ns indicates not significant by Mann-Whitney U test. e) Box plots showing total brood of wild-type animals and three isolates of animals lacking *sdg-2* (*Δ1, Δ2*, and *Δ3*) (top), and of animals with endogenous *sdg-2* tagged using *mCherryΔpi* (*sdg-2::mCherryΔpi*) along with paired wild-type animals (bottom). See Supplementary Fig. 9 for variation in *sdg-2(Δ1)*. f) Representative image of one gonad arm showing expression of SDG-2::mCherry fusion protein (left) and transcriptional reporter of *sdg-2* (right), with zoom of spermatheca (red box). Numbers of gonad arms imaged (n) and percentages showing the expression pattern are indicated. Scale bars, 20µm. h) Representative images of one gonad arm showing expression of SDG-2::mCherry (black or magenta) and CYLC-2::mNeonGreen (CYLC-2::mNG; black or green), with zoom of region showing proximal germ cells during early (top) and late (bottom) stages of spermatogenesis (red box). Percentages and n are as in (b). Scale bars, 20µm. Asterisks, ns, and n in (c) to (e) are as in Fig. 4b.

Taken together, these results suggest that SDG-1::mCherry made from the *sdg-1::mCherryΔpi* fusion transcript, which is ~16x more abundant than *sdg-1* mRNA, can cause a delay in oogenesis with a reduction in early progeny and that some lineages of animals lacking *sdg-1* could have a reduced brood size.

### SDG-2 shares sequence similarity with TRA-2 and is also predicted to interact with TRA-1

SDG-2 is a 77 amino-acid protein with sequence similarity to a region of Transformer (TRA-2) (Fig. 5a; Supplementary Fig. 7), which regulates gametogenesis. Specifically, TRA-2 interacts with the Gli-type transcription factor TRA-1 and the interaction promotes spermatogenesis (Kuwabara et al. 1998; Lum et al. 2000; Wang and Kimble 2001). This region of TRA-2, which is also present in its shortest predicted isoform TRA-2c, was reported to share similarity with two other known paralogs that were named XTR-1 and XTR-2 (Kuwabara et al. 1998). A BLAST search supports SDG-2 as another protein that shares sequence similarity, with particularly high identity in the C-terminal half of SDG-2, which we refer to as the SDG-2 TRA-2 homology (STH) region (Fig. 5a). This region in TRA-2 is also C-terminal after its MX region, which has a concentration of missense mutations in *tra-2* that were identified through genetic screens and that disrupt the TRA-2:TRA-1 interaction (Kuwabara et al. 1998; Lum et al. 2000; Wang and Kimble 2001). However, biochemical characterization using a truncation series had only reduced the potential interacting domains to larger regions: 1322aa-1475aa of TRA-2 ((Wang and Kimble 2001); corresponding to 234aa-387aa of TRA-2c) and 647aa-1110aa of TRA-1 (Lum et al. 2000). While the structures of TRA-2c and of SDG-2 are not predicted by AlphaFold3 with high confidence, the STH region in both proteins is predicted to be similarly alpha helical (Fig. 5b) and high-confidence predictions (Predicted Aligned Error (PAE) < 5Å and distance < 6Å) support an SDG-2:TRA-1 interaction (Supplementary Fig. 8, bottom right; Fig. 5c). Aligning the sequences of TRA-2c and all three related proteins (Fig. 5d; Supplementary Fig. 7a) revealed that they share different extents of similarity within the MX and STH regions (Fig. 5d; Supplementary Fig. 7b). XTR-1 has both parts with 76.2% identity within the MX region but only 37% identity within the STH region. XTR-2 is truncated after its MX region, which has 50% identity. SDG-2 has both parts with a limited identity of 18.2% within the MX region, but with 60.9% identity within the STH region. Consistently, AlphaFold3 also predicts interactions of high confidence between these fragments (Supplementary Fig. 8, top left). Using AlphaFold3 to predict possible interactions of XTR-1, XTR-2, or SDG-2 with 647aa-1110aa of TRA-1 reveal confident predictions of interactions for all pairs. For XTR-1, both MX and STH regions are predicted to be constrained by inter-protein interactions (Supplementary Fig. 8, top right). For XTR-2, which only has the MX domain, an interaction via the MX region is predicted (Supplementary Fig. 8, bottom left). For SDG-2, an interaction via the STH domain is predicted (Supplementary Fig. 8, bottom right). Therefore, these results suggest that despite the concentration of missense mutations in the MX region (Kuwabara et al. 1998; Lum et al. 2000; Wang and Kimble 2001), both MX and STH regions are likely important for interactions with TRA-1 in the case of TRA-2 and other proteins with sequence similarity, including SDG-2 (Fig. 5e).

**Fig. 7.**
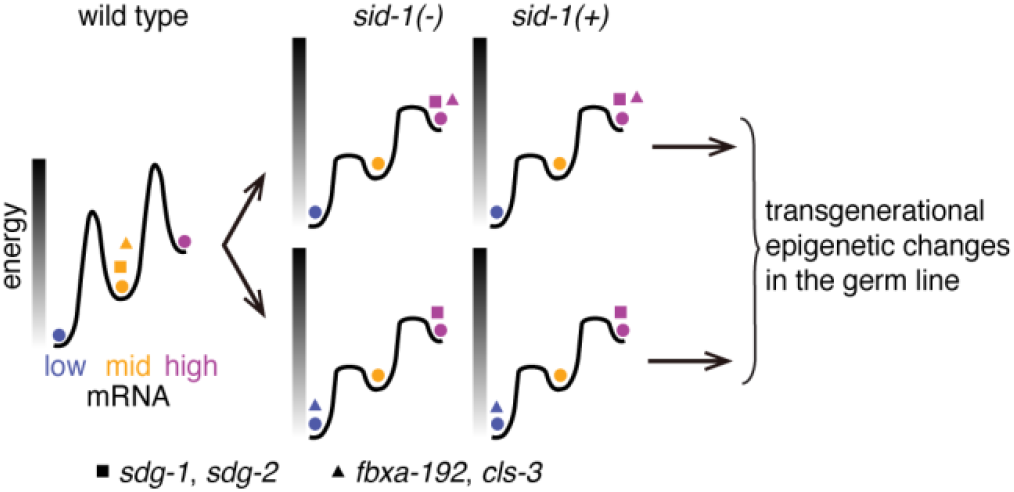
Conceptual model for transgenerational changes in SID-1-dependent genes. Schematic illustrating mRNA levels of different genes (shapes) that can either increase (high, magenta) or decrease (low, blue) from their prior levels (mid, orange) when *sid-1* is perturbed, ultimately causing transgenerational epigenetic changes in the germ line. The high energy (black) required for switching expression states in wild-type animals is depicted as barriers between levels. Changes in the abundance of *sdg* mRNAs suggest that the barrier between heritable expression states is reduced in *sid-1(−)* animals and can remain reduced upon reversion (*sid-1(+)*). Some changes (e.g., *sdg-1* and *sdg-2*, square) are consistent across generations, suggesting the stable maintenance of an altered state. Other changes (e.g., *fbxa-192* and *cls-3*, triangle) can vary between populations and yet appear to be maintained at high (top row) or low (bottom row) levels across generations. Genes that maintain their relative mRNA abundance in all cases (circles) are also indicated for reference. See text for details.

### SDG-2::mCherry is expressed in the germ line and is overexpressed upon loss of SID-1

To visualize where the SDG-2 protein might be present, we tagged the endogenous *sdg-2* gene with *mCherryΔpi* (Fig. 6a) and examined fluorescence from the resultant SDG-2::mCherry fusion protein (Fig. 6b). Uniform fluorescence from SDG-2::mCherry was observed in the hermaphrodite germ line and in the embryos held *in utero* (Fig. 6b), with a notable depletion in sperm. This SDG-2::mCherry fluorescence was also increased by ~1.4-fold in animals with a premature stop codon in *sid-1* (*sid-1(non)*) and after reversion to wild type (*sid-1(rev)*) (Fig. 6c), suggesting that the SID-1-dependent transgenerational changes observed for *sdg-2* mRNA (Fig. 1c; (Shugarts Devanapally et al. 2025)) lead to similar persistent changes in protein levels. Such *sid-1*-dependent changes were not observed for mCherry expression from a transcriptional reporter (Fig. 6d; see Fig. 6a for schematic).

### Loss of SDG-2 reduces brood size in some isolates

To evaluate the role of *sdg-2*, we generated three isolates with the *sdg-2* ORF deleted using genome editing (Fig. 6a). These isolates showed differences in brood size: one had a normal median brood size (*sdg-2(Δ1)* in Fig. 6e), another had an outlier with a low brood but an otherwise normal median brood size (*sdg-2(Δ2)* in Fig. 6e), and a third had an overall significantly lower median brood than that of wild-type animals (*sdg-2(Δ3)* in Fig. 6e). Consistent with the normal brood size of *sdg-2(Δ1)* animals, the numbers of sperm counted after DAPI staining of *sdg-2(Δ1)* animals were comparable to those similarly counted in wild-type animals (Supplementary Fig. 9a-b). However, when another cohort of *sdg-2(Δ1)* animals were measured at a different generation, these animals had a significantly lower brood size than that of wild-type animals (Supplementary Fig. 9c), indicating that loss of *sdg-2* can have a varying effect on brood size across isolates and across generations of the same isolate.

Since a sperm defect can cause a reduction in brood size (Ward and Carrel 1979) and since SDG-2 is predicted to interact TRA-1 (Fig. 5c), potentially regulating sperm production, we examined in greater detail the expression of SDG-2::mCherry in the spermatogenic germ line and in sperm. No defect in brood size was detected in animals with the endogenous *mCherryΔpi* tag of *sdg-2* (Fig. 6e, bottom), suggesting that tagging SDG-2 does not impact germline function. Unlike the bright SDG-2::mCherry expression observed in the oogenic germ line, SDG-2::mCherry was only faintly detected in sperm (Fig. 6f, left). In support of this faint expression being signal from SDG-2::mCherry, the transcriptional reporter driving mCherry under the control of *sdg-2 cis*-regulatory sequences showed bright expression within the sperm (Fig. 6f, right). Using L4-staged worms, we found that SDG-2::mCherry is present in the spermatogenic germ line (Fig. 6g) and like SDG-1::mCherry (Supplementary Fig. 4a, bottom) accumulates in CYLC-2-lacking structures (which are likely residual bodies) as part of the progression from spermatocytes to spermatids (Fig. 6g), explaining the low levels of SDG-2::mCherry in sperm (Fig. 6f). Given the presence of SDG-2::mCherry in the spermatogenic germ line of the hermaphrodite, the brood size defect seen in some populations of *sdg-2(Δ)* animals (*sdg-2(Δ3)* animals in Fig. 6e and *sdg-2(Δ1)* animals in Supplementary Fig. 9c) merits further investigation.

### Discussion

We have expanded the list of possible SID-1-dependent genes (SDGs) and analyzed two SDGs in detail to reveal processes under SID-1-dependent regulation. The patterns of changes observed for some *sdg* mRNAs suggest that loss of SID-1 causes a transgenerational change in the barrier between expression states for these genes (Fig. 7). Despite the *sdg-1* gene being located within a retrotransposon, the SDG-1 protein is expressed within the germ line and in embryos, with conditional enrichment in nuclei, near perinuclear granules, and at microtubules. Animals with *sdg-1::mCherryΔpi* mRNA, which is overexpressed when compared with *sdg-1* mRNA, have germline defects and altered dynamics of progeny production. SDG-2 has the potential to interact with TRA-1 via its STH domain to influence spermatogenesis and/or sperm function. However, only some lineages of animals lacking *sdg-1* or *sdg-2* show a reduced brood size. Thus, changes in the expression levels of two different SID-1-dependent genes can alter *C. elegans* reproduction and could be regulated to impact organismal fitness.

### Processes regulated by SID-1

Both the relevant molecular function(s) of SID-1 and the different regulatory networks that each SDG integrates into need to be understood to establish the process(es) regulated by SID-1. The evolved molecular function(s) of SID-1 *in vivo* are unknown. While dsRNA-binding is well-supported as one function of SID-1 proteins by Cryo-EM structures (Wang et al. 2024; Zhang et al. 2024) and by the requirement of SID-1 for dsRNA entry into the cytosol (Winston et al. 2002; Feinberg and Hunter 2003), analysis of SID-1 and/or its mammalian homologs raise the possibility of additional functions such as binding or cleaving lipids (Qian et al. 2023; Sun et al. 2023b; Hirano et al. 2024; Liu et al. 2024a; Navratna et al. 2024; Zhang et al. 2024; Zheng et al. 2024). Furthermore, for any protein, signaling through protein-protein interactions rather than intrinsic biochemical activity cannot be excluded as a key function. Therefore, changes in any of the 157 protein-coding genes or pseudogenes identified here as possibly SID-1-dependent (Fig. 1a) need not be caused exclusively by changes in the import of dsRNA. While we could search for unifying themes for all SID-1-dependent genes, the analysis of *sdg-1* and *sdg-2* presented here already suggest that multiple processes within the germ line are perturbed by changes in *sid-1*. The magnitude of changes in mRNA levels of many SDGs detected using RNA-seq is less than 2-fold: 47 of 96 changed in *sid-1(deletion)*, 10 of 12 changed in *sid-1(nonsense)*, and 54 of 70 in *sid-1(reversion)*. These modest changes could reflect minimal impact of SID-1-dependent regulation throughout the organism or alternatively conditional (e.g., stage-specific or tissue-specific) regulation of higher magnitude that is diluted when total RNA from mixed-stage populations is examined. Given the dynamic expression pattern of SID-1 throughout development, from ubiquitous in hatched L1-staged larval animals to dramatic tissue-specific enrichments in later development (Shugarts Devanapally et al. 2025), additional analyses (e.g., tissue-specific, stage-specific, or environment-specific) are needed to determine when and where each SDG is regulated by SID-1. The *sid-1*-dependent initiation of persistent changes in *sdg-1* and *sdg-2* is reminiscent of *sid-1*-dependent enhanced initiation of the persistent silencing of a single-copy transgene upon mating (Shugarts Devanapally et al. 2025). Both *sdg-1* and *sdg-2* are targeted by piRNAs (multiple studies summarized in (Shugarts Devanapally et al. 2025)) and mating-induced silencing requires piRNA binding to the silenced gene (Devanapally et al. 2021). Consistent with interaction between different small RNA-mediated mechanisms within the germ line, piRNA-binding sites on a gene promote the persistent silencing of that gene by a homologous silenced gene (Devanapally et al. 2021) and loss of the piRNA-binding Argonaute PRG-1, which alters the expression of many genes (e.g., (Barucci et al. 2020; Reed et al. 2020)), can result in the transgenerational stability of silencing by extracellular dsRNA (Shukla et al. 2021) through mechanisms that remain unknown. Therefore, a reasonable hypothesis to pursue in future studies is that changes in SID-1 alter the unknown population of imported extracellular dsRNAs and these imported dsRNAs compete with piRNA-mediated regulation within the germ line to reduce transgenerational changes in SDGs. Since different SDGs could antagonize each other, detecting the phenotypic consequences of the simultaneous changes in many SDGs expected in *sid-1(−)* animals could require specific environmental conditions or multiple generations.

We found that *cls-3* and *fbxa-192*, identified here as *sid-1*-dependent genes, show transgenerational changes that do not always maintain the direction of change (Fig. 1c and Supplementary File 1). Our hypothesis that loss of *sid-1* results in unbuffered gene expression because the barrier for change is lowered for these genes (Fig. 7) accommodates this result as a specific consequence of SID-1 loss and thus potentially the loss of regulation by extracellular dsRNA. However, an alternative possibility is that such genes simply have mRNA levels that vary widely across cohorts of wild-type animals. Under this hypothesis, their mRNA levels will have a strong dependence on the founding populations being cultured. As a result, when any mutant lacking gene-*x* is compared with ‘wild type’ through an RNA-seq experiment, genes with unstable expression that nevertheless can persist for many generations could be identified as an *x*-dependent gene based on the cohorts of animals sequenced. These genes with unstable expression would be among the genes present in multiple RNA-seq datasets generated in the field of RNA silencing in *C. elegans*, along with genes that are frequently perturbed. In a recent study (Lalit and Jose 2025), a list of the top 100 such frequently identified genes was aggregated and indeed includes some of the genes categorized as *sid-1*-dependent in this study. While the germline-expressed gene *fbxa-192* (Almeida et al. 2025) is in this top 100 list (34^th^ *r*_*100*_ score), *cls-3* is not. Therefore, future analyses are needed to distinguish between these two explanations for apparently *sid-1*-dependent genes that do not maintain their direction of change.

### Roles of SDG-1

SDG-1 is likely a multi-functional protein with dynamic localization and an impact on germline function. As a retrotransposon-resident gene, *sdg-1* is subject to transgenerational silencing by small RNAs (Rojas et al. 2025; Shugarts Devanapally et al. 2025) and yet it is expressed and translated into a protein (Fig. 2b). Although the predicted structure of the SDG-1 protein is not of high confidence (Fig. 2d), it weakly resembles a dsRNA-binding protein (Fig. 2f) and animals lacking *sdg-1* show a small enhancement of dsRNA-mediated gene silencing (Fig. 2h), suggesting a possible interaction of SDG-1 with dsRNA. Biochemical and structural studies are needed to evaluate this possibility. Beyond the molecular function(s) of SDG-1, the various subcellular localizations observed using the SDG-1::mCherry fusion protein (Fig. 3) suggest that SDG-1 could play a role in multiple processes. These localizations include enrichment within dividing nuclei but exclusion from non-dividing nuclei (Fig. 3, Supplementary Fig. 3; (Shugarts Devanapally et al. 2025)), conditional enrichment near germ granules (Fig. 3d), and conditional recruitment to microtubules (Fig. 3e). We speculate that the different localizations could be in response to different signals. Exclusion from the nucleus could be regulated by post-translational modifications (e.g., Thr5, Ser6, and Thr8 of SDG-1 are predicted to be phosphorylated (Chen et al. 2023)). Germline enrichment could be driven by the stress of imaging conditions (e.g., reduced oxygen, increased temperature, restricted movement, etc.). Recruitment to microtubules, which is only observed in ~66% of animals that show germ granule enrichment, could be a response to prolonged sensing of compressive forces when the worm is sandwiched between a slide and a coverslip (Supplementary File 4). One of only four other SID-1-dependent genes detected in all three comparisons with wild-type animals (Fig. 1a) is *cls-3*, which encodes a homolog of CLASP proteins that are recruited for microtubule repair and stabilization in response to compressive forces (Lawrence et al. 2020; Ju et al. 2024). Consistently, CLS-3 and its paralogs have a role in the assembly of astral microtubules in early embryos (Espiritu et al. 2012). The SDG-1::mCherry expressed from an endogenous tag generated using genome editing resulted in morphological defects within the germ line and in animals that laid fewer early progeny but ultimately had a normal brood size (Fig. 4). A unifying speculation inspired by the nuclear and microtubule localizations of SDG-1 (Fig. 3) and the dynamics of germline defects in *sdg-1(−)* animals (Fig. 4) is that SDG-1 overexpression results in prolonged mitosis, which delays subsequent meiosis and ovulation, detected as fewer early progeny but more later progeny, yielding a nearly wild-type brood. Determining the physiologically important signals that could be driving each of the localizations of SDG-1 and mapping each localization to defect(s) in the germ line requires more experiments.

### Roles of SDG-2

SDG-2 is a small protein (Fig. 5) that is predicted to interact with the Gli-type transcription factor TRA-1 (Zarkower and Hodgkin 1992) and like the TRA-2:TRA-1 interaction (Lum et al. 2000; Wang and Kimble 2001), the SDG-2:TRA-1 interaction could promote sperm production and/or function. The primary sequence of SDG-2 has a 38 amino-acid region that is >60% identical to a region of TRA-2 (Fig. 5a) and is part of the region that was experimentally shown to interact with TRA-1 (Lum et al. 2000; Wang and Kimble 2001). This STH region is also present in another paralog of TRA-2 called XTR-1 but not in XTR-2. Together, these interactors of TRA-1 could all be promoting spermatogenesis or could be engaged in more complex regulation where the activity of TRA-1 is responsive to environmental or developmental signals relayed by these TRA-1-interacting proteins, including SDG-2. In support of the possibility of complex regulatory outcomes for the TRA-1:TRA-2 interaction despite conservation of this interaction in related nematodes, identical mutations in the interacting domain of TRA-2 promotes oogenesis in *C. elegans* but promote spermatogenesis in *C. briggsae* (Shen et al. 2024). Consistent with responsiveness to environmental signals, the *sdg-2* gene was recently reported to show >4-fold upregulation of mRNA with >4-fold downregulation of antisense small RNA upon loss of *tcer-1*, a gene that suppresses immune responses against pathogenic bacteria (Y102A5C.36 in (Naim et al. 2026)). Alternatively, a constitutive role for *sdg-2* is suggested by the reduction in brood size observed in *sdg-2(Δ3)* animals (Fig. 6e) and in some cohorts of *sdg-2(Δ1)* animals (Supplementary Fig. 9), which could be the result of enhanced epigenetic variation in phenotype among sibling lineages lacking *sdg-2*. Determining how SDG-2 impacts gene regulation by TRA-1 and brood size requires additional experiments.

## Acknowledgements

We thank Maria Johnsonbaugh for early brood size assays evaluating *sdg-2(−)* animals and Kevin O’Connell, Eric Haag, and members of the Jose lab for comments on the manuscript. Some strains were provided by the CGC, which is funded by NIH Office of Research Infrastructure Programs (P40 OD010440). This work was supported by UMD CMNS Dean’s Matching Award for “Training Program in Cell and Molecular Biology” T32GM080201 to NMSD and in part by National Institutes of Health Grant R01GM124356 to AMJ.

## Notes

**Competing Interest Statement:** The authors declare no conflict of interest.

https://github.com/AntonyJose-Lab/Sathya_et_al_2026

https://www.ncbi.nlm.nih.gov/sra/PRJNA1475530

### Competing Interest Statement

The authors have declared no competing interest.

