## Supplementary Material for "Two SID-1-dependent genes sensitive to heritable epigenetic changes can also impact reproduction"

*Corresponding author: Antony M. Jose

**Competing Interest Statement:** The authors declare no conflict of interest.

**Keywords:** RNAi, transgenerational epigenetic inheritance, germline, sperm

**Running head:** SID-1-dependent genes regulate fertility

This supplement has the following:

2 Supplementary Tables

9 Supplementary Figures

4 Supplementary File titles

**Supplementary Tables**

**Table 1.** Strains used.

| Name | Genotype |
|---|---|
| N2 | wild type |
| AMJ1159 | *sid-1(jam80[nonsense]) V* |
| AMJ1170 | *jamSi37 [mex-5p::mCherry::cye-1 3'UTR + unc-119(+)] II; unc-119(ed3) III* |
| AMJ1217 | *sid-1(jam86[revertant]) V* |
| AMJ1324 | *sid-1(jam113[deletion]) V* |
| AMJ1347 | *Y102A5C.36(jam126[deletion]) V* (*sdg-2(Δ1)* in paper) |
| AMJ1348 | *Y102A5C.36(jam127[deletion]) V* (*sdg-2(Δ2)* in paper) |
| AMJ1349 | *Y102A5C.36(jam128[deletion]) V* (*sdg-2(Δ3)* in paper) |
| AMJ1371 | *Y102A5C.36(jam136[Y102A5C.36::mCherryΔpi]) V* |
| AMJ1372 | *W09B7.2(jam137[W09B7.2::mCherryΔpi]) F07B7.2(jam137[F07B7.2::mCherryΔpi]) V* |
| AMJ1388 | *Y102A5C.36(jam136[Y102A5C.36::mCherryΔpi]) sid-1(jam149) V* |
| AMJ1389 | *W09B7.2(jam137[W09B7.2::mCherryΔpi]) F07B7.2(jam137[F07B7.2::mCherryΔpi]) sid-1(jam150[nonsense]) V* |
| AMJ1399 | *sid-1(jam157[nonsense]) V* |
| AMJ1405 | *sid-1(jam163[revertant]) V* |
| AMJ1406 | *sid-1(jam164[revertant]) V* |
| AMJ1407 | *sid-1(jam165[revertant]) V* |
| AMJ1411 | *Y102A5C.36(jam136[Y102A5C.36::mCherryΔpi]) sid-1(jam169) V* |
| AMJ1446 | *W09B7.2(jam137[W09B7.2::mCherryΔpi]) F07B7.2(jam137[F07B7.2::mCherryΔpi]) sid-1(jam177[nonsense]) V* |
| AMJ1486 | *Y102A5C.36(jamSi68[y102a5c.36p::mCherryΔpi]) V* |
| AMJ1576 | *W09B7.2(jam231[deletion]) F07B7.2(jam231[deletion]) V* (*sdg-1(Δ3)* in paper) |
| AMJ1577 | *W09B7.2(jam232[deletion]) F07B7.2(jam232[deletion]) V* (*sdg-1(Δ1 or Δ)* in paper) |
| AMJ1612 | *W09B7.2(jam241[deletion]) F07B7.2(jam241[deletion]) V* (*sdg-1(Δ2)* in paper) |
| AMJ1615 | *W09B7.2(jam244[sdg-1 ΔORF in jam137]) F07B7.2(jam244[sdg-1 ΔORF in jam137]) V* |
| AMJ1860 | *adr-2(uu28) III* |
| AMJ1862 | *adr-2(uu28) III; W09B7.2(jam232[deletion]) F07B7.2(jam232[deletion]) V* |
| AMJ1863 | *adr-2(uu28) III; W09B7.2(jam232[deletion]) F07B7.2(jam232[deletion]) V* |
| AMJ2006 | *cylc-2(mon2[cylc-2::mNG::3xFLAG) I; W09B7.2(jam137[W09B7.2::mCherryΔpi]) F07B7.2(jam137[F07B7.2::mCherryΔpi]) V* |
| AMJ2077 | *W09B7.2(jam241[deletion]) F07B7.2(jam241[deletion]) V* (2x outcross isolate #2) |
| AMJ2078 | *W09B7.2(jam241[deletion]) F07B7.2(jam241[deletion]) V* (2x outcross isolate #1) |
| AMJ2084 | *W09B7.2(jam137[W09B7.2::mCherryΔpi]) F07B7.2(jam137[F07B7.2::mCherryΔpi]) V* (2x outcross, isolate #2) |
| AMJ2085 | *W09B7.2(jam137[W09B7.2::mCherryΔpi]) F07B7.2(jam137[F07B7.2::mCherryΔpi]) V* (2x outcross, isolate #1) |
| AMJ2109 | *cylc-2(mon2[cylc-2::mNG::3xFLAG) I; Y102A5C.36(jam136[Y102A5C.36::mCherryΔpi]) V* |
| BB239 | *adr-1(uu49) I; adr-2(uu28) III* |
| MDX44 | *cylc-2(mon2[cylc-2::mNG::3xFLAG) I* |
| WM49 | *rde-4(ne301) III* |

**Table 2.** Oligonucleotides used.

| Name | Sequence (5'>3') |
|---|---|
| P1 | cgtggcacatactttccgttgttg |
| P2 | tgcacggcgtatcaaactg |
| P3 | ccgcaagtctctcctgtatg |
| P4 | gctgctcaagcaaatcgaatg |
| P5 | agggagtatgtctgtgagcttg |
| P6 | cagtacaacacgagcctttgg |
| P7 | gcgtatttgaagctcggctc |
| P8 | gcgtcttcttcgtgatcttcg |
| P9 | aatgtttccgaccgcagc |
| P10 | gtcatctccgacgagcac |
| P11 | ttccgttgttggcttcgttg |
| P12 | ggccattgggagaacttcg |
| P13 | tgacggcctcttctacatatcg |
| P14 | gctgaaggtggatagtgtctc |
| P15 | attgctccgcaaatgtagtgg |
| P16 | ttatcacggtggagaacagc |
| P17 | ttggtagggaatcggctgg |
| P18 | tccacgtcatcattcagcttg |
| P19 | cgagttcttcactgcatctgac |
| P20 | cgctgaaactctgcccaac |
| P21 | gtatcggatcgcacattagctc |
| P22 | cgacggggattgacaacttt |
| P23 | gaagctcggctctttcatg |
| P24 | tgcggcgagaaagctgatg |
| P25 | ggcaatgctcaagcttctga |
| P26 | ccgcaaggacattgacagc |
| P27 | aggctcaacagactcgtacttc |
| P28 | acgattggttttcttccatg |
| P29 | accttttcctcaagtgtgtg |
| P30 | caggatcgtcgcaaagagtg |
| P31 | ctcattccttgtgcaagctg |
| P32 | cggttgatgttcctcggatg |
| P33 | gtgatctctcccaagtgttg |

| | |
|---|---|
| P34 | attgcatgttgctgcacatc |
| P35 | atcgaggagacactatccac |
| P36 | aatgtagtggagcgatgctg |
| P37 | atgtgagcgggatttggttc |
| P38 | gtacgggtactgatgcatgg |
| P39 | atcatgtagagcggttggtg |
| P40 | atcgtggagagttgtgcaag |
| P41 | tcgtgttccgtcgatcaaga |
| P42 | tgcatttcctcaaacgggac |
| P43 | actggaggtcgataacgagc |
| P44 | gttcactagtcgatgttgctc |
| P45 | aatcacatgggtcactgatgc |
| P46 | acagtttccttcacaaagtcg |
| P47 | gtgtctacttaagaacgtggag |
| P48 | cgaaagcagcaagagtgaagg |
| P49 | atcagtgccagagccatag |
| P50 | gagttcggaagtaaaccgtgg |
| P51 | gctgaaggtggatagtgtctc |
| P52 | cgcagtacgcagagtgaac |
| P53 | gagttcggaagtaaaccgtgg |
| P54 | tctcccacttgaatccctctg |
| P55 | gctgaaggtggatagtgtctc |
| P56 | cgacatcgtggaaaccgac |
| P57 | cttcatacactgctccgctg |
| P58 | cgacatcgtggaaaccgac |
| P59 | cttcatacactgctccgctg |
| P60 | ttggtagggaatcggctgg |
| P61 | caccttcgccaattatcacctc |
| P62 | cgtcagcttctgattcgacaac |
| P63 | ucccuccaaacagacaaugaguuuuagagcuaugcu |
| P64 | gaaggcgauuguucaguuccguuuuagagcuaugcu |
| P65 | ttttttgaattcatttaacatttccagaaagccgtggacgacatcgtggaaaccgactcgaatc aactatccgatcaaagacattccctccaaacagacaattttatcttaccaatttttgttgttaat tttttatagataaacaataaaaatgctgaaaaatttgatcatttgaaaatcttgaaatttcaatt ggaga |
| P66 | gaaggcgauuguucaguuccguuuuagagcuaugcu |
| P67 | agcgagatgaaggcgattgctcggtcccaggtgttatggtctccaagggagagga |
| P68 | aaaattaacaacaaaaattggtaagataaaatttagccactacctgatcccttgt |

| | |
|---|---|
| P69 | ctagttctagacattctctaatgaaaaaatctttcag |
| P70 | aagtgcaagtaagatcagtgtttgtttc |
| P71 | actgatcttacttgcacttatgaaaaatgattaaataattaaatggataatgcacaaaga |
| P72 | tggagaccattgtctgtttggagggaatgtct |
| P73 | caaacagacaatggtctccaagggagagga |
| P74 | taaaatttagccactacctgatcccttgt |
| P75 | aggtagtggctaaattttatcttaccaatttttgttgttaatttttttatagataaaca |
| P76 | tagagaatgtctagaactaggtcatttttttgggaaattagatgaagagc |
| P77 | agcauucaaucgagacugcaguuuuagagcuaugcu |
| P78 | agcctataatctatatcagcattcaatcaaggctacacggttacgatcaggttttgatggaaat gagggt |
| P79 | agcauucaaucgagacugcaguuuuagagcuaugcu |
| P80 | aagcctataatctatatcagcattcaatcgagactgcacggttacgatcaggttttgatggaa atgaggg |
| P81 | tcaggagaagaaggacgatg |
| P82 | tttccgatgactgtgactgttc |
| P83 | gtggacttgaggttgagct |

**Supplementary Figures**

a

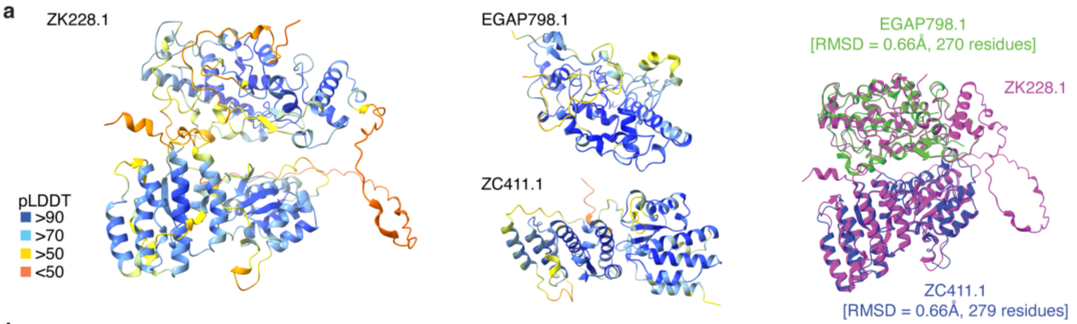


b

```
  ZK228.1   1 MEDNPDYRPHEVERYPVDLPRQHPRPGILDIRCEFTSIKEFFGEEPPFNERTVLRFAEYVRGYYPDHQPTHVMDRGGLALYIWIDRMPTPATHLQQTFLRL
EGAP798.1   1 -MGDPDERPWEIERVPIQLPRQHPAPIVEDIRYEALNIFQFFGIRFPFDEQTIIRFATLEENFPPEFKPNNVMERGGILLFIWADRLPTPATHLQQSIIRL
  ZC411.1     -----------------------------------------------------------------------------------------------------

  ZK228.1 102 PFLEEVENLSRFVLRHARAQNEHLEGERTNRCSVCGSHLLGMPLADHSYIRCPLDALEGERRLEFLAINHAAFCSKCNSRFAKHTGCTGPVCHSCDQRGHT
EGAP798.1 101 PLLGQIADLPRFVFLYARAQNEPREGDRTNRCPLCGSHLLGLPLADHRLENCPFGILEGEQRLEFLAINHVSYCHMCNSRSATHSQCARPPCRQCNIVGHT
  ZC411.1     -----------------------------------------------------------------------------------------------------

  ZK228.1 203 AATRLCDSRLGRLTPESQQELLTVVTQHRRNYLLHIRNLTNDINKPLRYRSYTDQAYIFLERRVEDR-WLIDGWGFYHDDAGEFPQPAMQRYSNH-PHRSV
EGAP798.1 202 VASGLCESPMGTTNLGQQHLLQQRVTTFRRNHLLRIRALTNRQDRPLQYRSYTDQAFIYLERRVQNRRNEMIGWSFYHDDAGEFPQHHPGRYSNYLEERPV
  ZC411.1     -----------------------------------------------------------------------------------------------------

  ZK228.1 302 RYKGLVPPEYHSQAVNPIPTFPPHEVEYLQAVHEVIIEFHRSGELHMERILPNAAIADEPVAIPVEVDPPPAVIVEEGIAIHPVENVDPLEMPVNEIHERN
EGAP798.1 303 HYLSLAPPEYHSQAVNRIPTFAPR-----------------------------------------------------------------------------
  ZC411.1     ----------------------------------------------------------------------------------------------------M

  ZK228.1 403 VADSEEDGDDDLILQDPEPLTVNNDTARPAFQHKAFETLLQHPLPEGKNALTLRVRALQVILCGRSGTEWEVSPEVSAALLRRYAKDLILYGQALKNEEGL
EGAP798.1     -----------------------------------------------------------------------------------------------------
  ZC411.1   2 I--TEARINGNAYINCQESLTLENDSARPAFQHRAFEGLIQHPLPLGQDALVSRVRALQVILCGRADPRYDAPFNGSPILLRQYAEDMIQYGHALKAEECL

  ZK228.1 504 IVQMEEFLSATVLRKERYGSDIVYIPSMSLWAKAEVRQSLTRLFDPHPFGDETTRRLDRVGHNHRYWFGLPSNQ-GALHEDLQVAEEERLSAQLVPTKARI
EGAP798.1     -----------------------------------------------------------------------------------------------------
  ZC411.1 101 MVQLEENEPNKVMRKQRYGADTVTLPTFDLWCIPEVRQSLYVLFQPHPFGDESTRCLDRFGNTNRYWFTLPISGEYTLLEQLQEDEERRLTFRFTPTNRCI

  ZK228.1 604 WESPLHDIGQLRNALEIPVPTDNQAVRLRIGWMIHLLTCQLPPETVNQGSYVEVLDYLKILRAATKLVLSIREEASTVRIELSTCMRHMATSSARGHNLVL
EGAP798.1     -----------------------------------------------------------------------------------------------------
  ZC411.1 202 WESPLHDVTKLRRALNTPLPTGQQALRLRLAWMINLLTGQKPPETVFQGTYVETRNFLKIVRGATMLVLGIREEASTVKVELSTCVRHIATSSARGHNLVL

  ZK228.1 705 PTFGLFVKYAMGKWESWLNGCWDHVYDEITLRHVESCPCHVDEAMFRYEY
EGAP798.1     --------------------------------------------------
  ZC411.1 303 PTFTLFTMYSMEKWEAWLNGCWNHIFNEITGRDIESCTCHQDEEME----
```

c

```
CERV-1 114 NRQLKEKSEELMQKSQILVETTLKLKAVEEERDKRKKEEQFREADARSNNYARKDHGYKMHNIELKNEYTSTTYRCRYICRCALKPCMFNMTLVPEAHTP
ERI-6c 354 NRHLNELRSDLREKKSVIFSQGLKIDMLESQLSQRQTEFEYRSANAAANSFTRKDHGKLYHCFDIKNQ-DSTSVSDTMTCRCSMKPCVFYLDSLPAAKTP

CERV-1 214 SPTQLYRMYCIMEKSGNRKIDPKQLMAMSSRPLPSPLQITLPDKMMDNLFKDMIGCSSLWTYVAELGWENSYNRYVDKLLNENCGDILNG
ERI-6c 453 SLTQLRRIVVLMQQKPSQKIDPKEMLLMTSMHLPFPMQ-----EPSTEYFNAISGCSTSYICVEEMGIGFSFRRHVEKLWMKNLPPATNG
```

**Suppl. Fig. 1.** Related to Fig. 2. Some retrotransposon-encoded proteins are highly similar to other proteins, suggesting recent evolution and/or selection for specific structure. a and b). EGAP798.1 and ZC411.1 have high structural (a) and sequence (b) similarity to the N-terminal and C-terminal halves of ZK228.1, respectively. Alignment of the structures predicted by Alphafold for ZK228.1, EGAP798.1 and ZC411.1 are shown (a, right). Root Mean Square Deviation (RMSD) and numbers of aligned residues are as indicated. Identical residues are indicated in purple (b). c) Part of the CERV-1 protein shares sequence homology with the RNA silencing protein ERI-6c. Identical residues are as in (b).

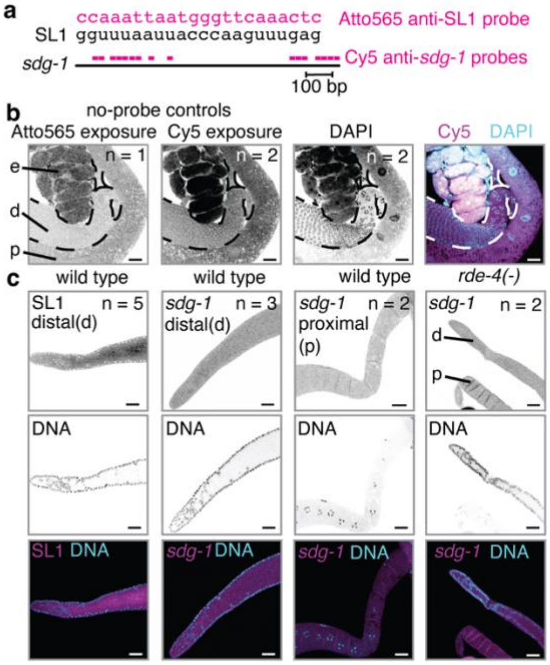


**Suppl. Fig. 2.** The *sdg-1* mRNA is not detectable above background using smFISH in wild-type animals and in *rde-4(-)* animals. a) Probes used. Top, Sequence of Atto565 labeled anti-SL1 probes (pink) complementary to SL1 sequence (black). Bottom, spliced *sdg-1* transcript targeted by a set of 16 Cy5 labeled anti-*sdg-1* probes (pink). b). Control imaging without probes and with DAPI staining of wild-type animals under conditions used for imaging Atto565 (first), Cy5 (second), and DAPI (third). A merge of DAPI with Cy5 is also shown (fourth). Numbers of dissected animals imaged (n), proximal and distal regions of gonads (p and d, respectively), and embryos (e) are highlighted. c) Top row, Representative images showing smFISH signal using anti-SL1 probe (column 1) or anti-*sdg-1* probe (columns 2-4) of dissected germlines from wild-type (columns 1-3) or *rde-4(-)* animals (column 4). Middle row, Corresponding DAPI staining. Bottom row, merge of smFISH signal (magenta) and DAPI (blue). Scale bars, 20 µm. Unlike the signal for SL1, the signal for *sdg-1* is not above background.

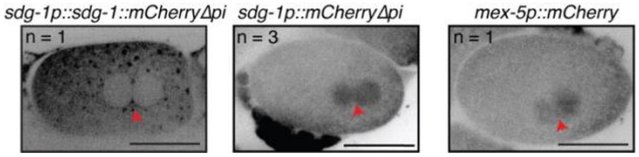


**Suppl. Fig. 3.** Representative images of pronuclei meeting in the 1-cell embryo. Unlike SDG-1::mCherry, mCherry alone remains enriched within nuclei before pronuclear fusion (red arrow) in 1-cell embryos (n indicates number of embryos imaged at this specific stage), although all were enriched within the nucleus in -1 oocytes. Scale bar, 20 µm.

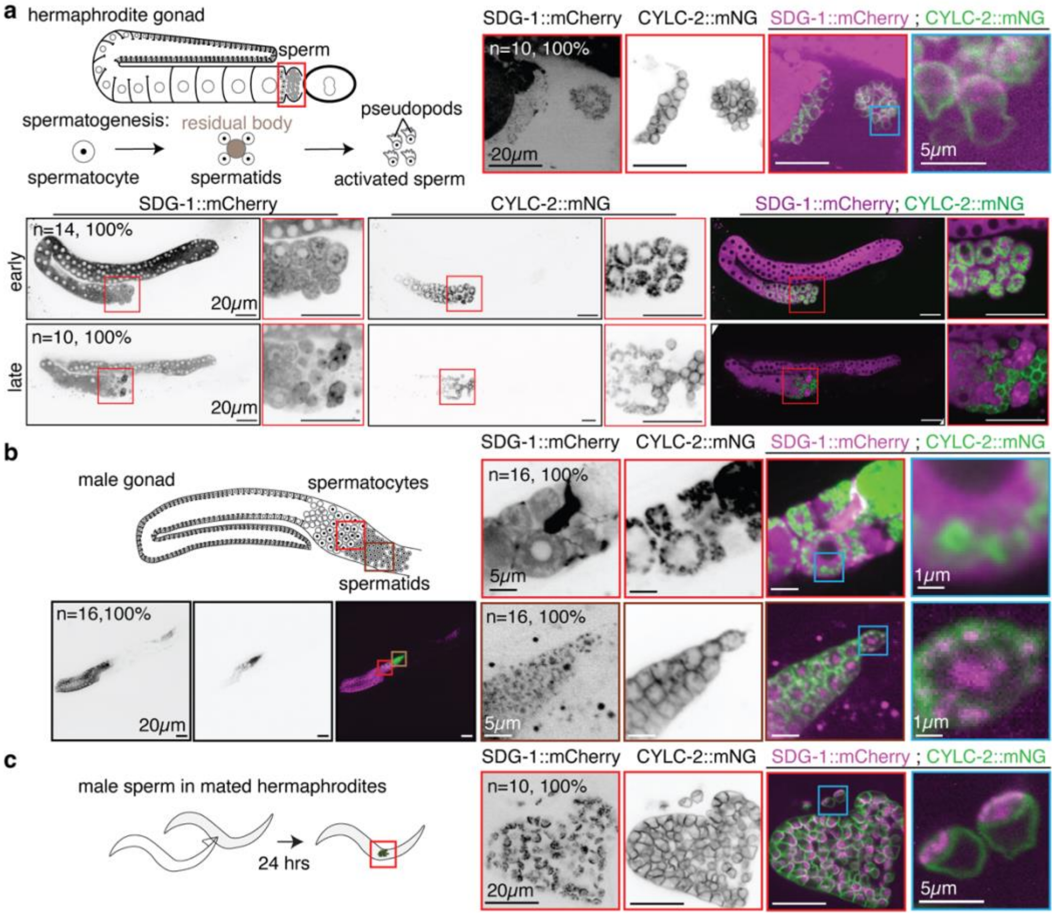


**Suppl. Fig. 4.** SDG-1 levels and subcellular localization are regulated during different stages of spermatogenesis. a) Top Row: Left, Schematic of hermaphrodite gonad arm and embryo with region where sperm is stored to be imaged (red box) shown above schematic of transformation from spermatocyte to spermatids along with residual body formation, followed by activation. Right, Representative images of spermatheca region showing expression of SDG-1::mCherry (black or magenta) and CYLC-2::mNeonGreen (CYCL-2::mNG; black or green), with zoom of region showing individual sperm (blue box). Numbers of spermatheca imaged (n) and percentages showing the expression pattern are indicated. Middle and Bottom rows: Representative images of one gonad arm showing expression of SDG-1::mCherry (black or magenta) and CYLC-2::mNeonGreen (CYCL-2::mNG; black or green), with zoom of region showing proximal germ cells during early (middle) and late (bottom) stages of spermatogenesis (red box). Number of gonad arms imaged (n) and percentages showing expression pattern are indicated. b) Top Left, Schematic of male gonad with regions of spermatocyte development to be imaged (red and brown boxes). Bottom Left, Representative images of male gonad showing expression of SDG-1::mCherry (black or magenta) and CYLC-2::mNeonGreen (CYCL-2::mNG; black or green) with merged image. Top Right, Zoom of red box from Bottom Left, including close-up view of part of a spermatocyte (blue box). Bottom Right: Zoom of brown box from Bottom Left, including close-up view of spermatid (blue box). Percentages and n are as in (a). c) In male sperm within the hermaphrodite germline SDG-1 localizes away from pseudopods. Left, Schematic showing mating and imaging of male sperm within hermaphrodite (red box). Right, Representative images of spermatheca region showing expression of SDG-1::mCherry (black or magenta) and CYLC-2::mNeonGreen (CYLC-2::mNG; black or green), with zoom of region showing individual sperm (blue box). Percentages and n are as in (a). Scale bars are as indicated.

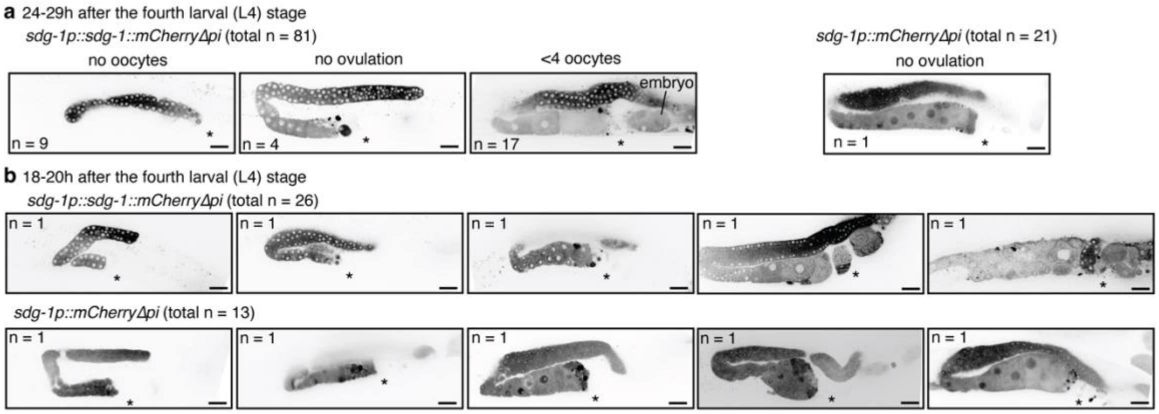


**Suppl. Fig. 5.** Related to Fig. 4. Gallery of defective germlines observed in animals with *sdg-1p::sdg-1::mCherry∆pi* or *sdg-1p::mCherry∆pi*. Representative images of a gonad arm showing defects as categorized in Fig. 4a (a) or additionally observed in a different cohort of animals (b) at a different time-point (18-20 h) after the L4-stage (determined using stage of vulval morphogenesis). Asterisks indicate location of the spermatheca. Scale bar, 20 µm.

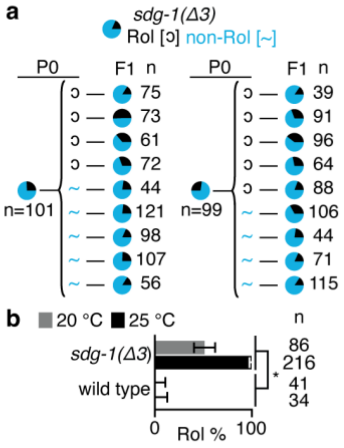


**Suppl. Fig. 6.** Related to Fig. 4. One isolate obtained after removing the *sdg-1* gene using genome editing shows a transmissible Rol defect that is enhanced at 25ºC. a) Pedigree charts of two experiments showing inheritance of Rol defect from parent to progeny in animals with a deletion in *sdg-1* (*sdg-1(∆3)*). Each pie chart shows the distribution of rolling (black) and non-rolling (blue) animals on the plate, with different total populations (n). Some progeny show a Rol defect (black), despite lack of a detectable defect in their parents (blue). b) Temperature dependence (20ºC, grey, versus 25ºC, black) of the Rol defect in wild-type and *sdg-1(∆3)* animals. Asterisks, error bars, and n are as in Fig. 2h.

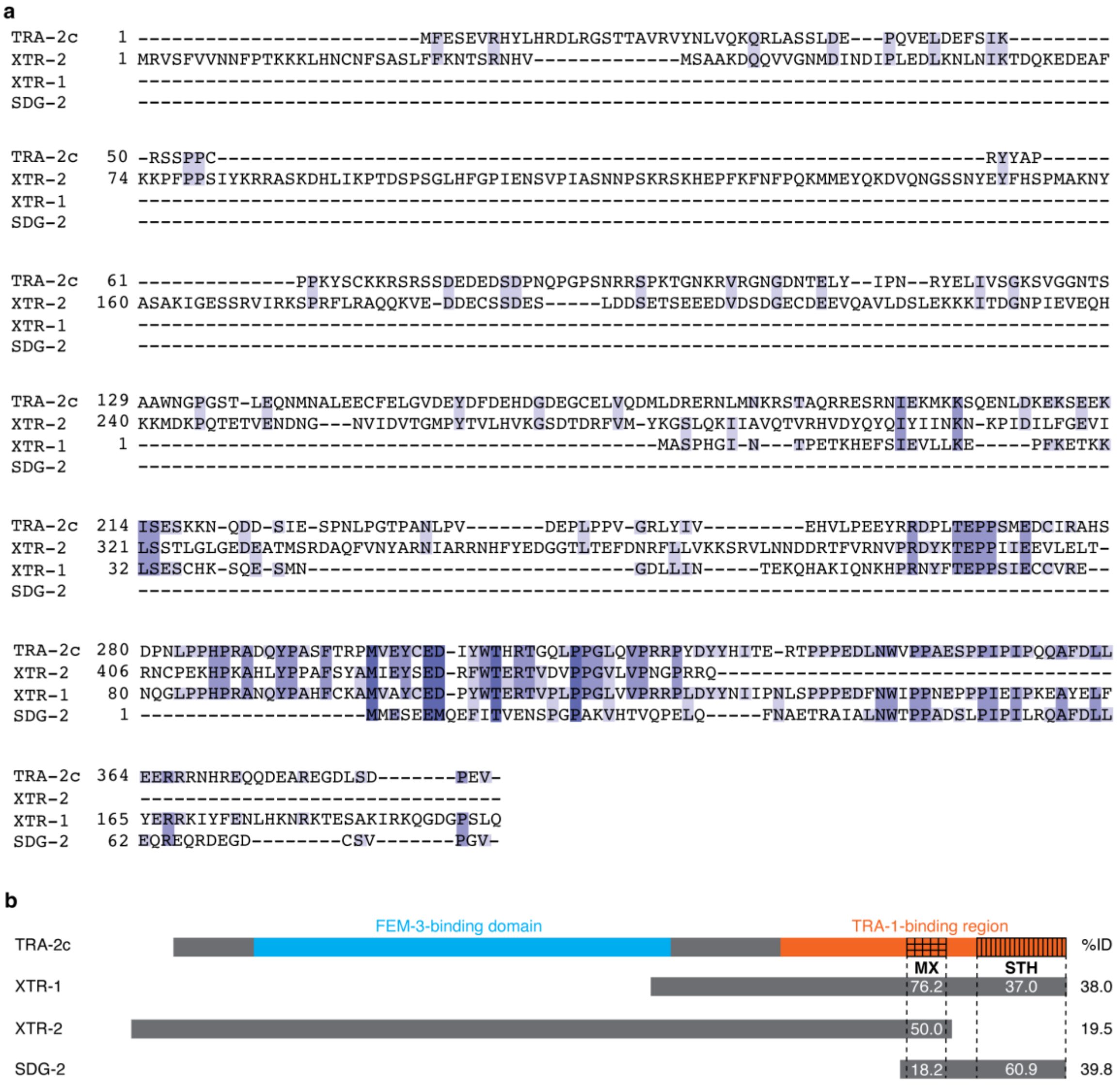


**Suppl. Fig. 7.** Related to Fig. 5. Protein sequence similarity between TRA-2c and related proteins, including SDG-2. a) The C-terminal region of TRA-2c shares sequence homology with three related proteins (XTR-1, XTR-2 (isoform a), and SDG-2). Residues are highlighted in shades of purple based on degree of conservation. b) Summary of alignment showing FEM-3 binding region and TRA-1-binding region within TRA-2c, which includes the MX and STH regions. The percentages of sequence identity overall (%ID) and within the MX regions and/or STH regions are indicated.

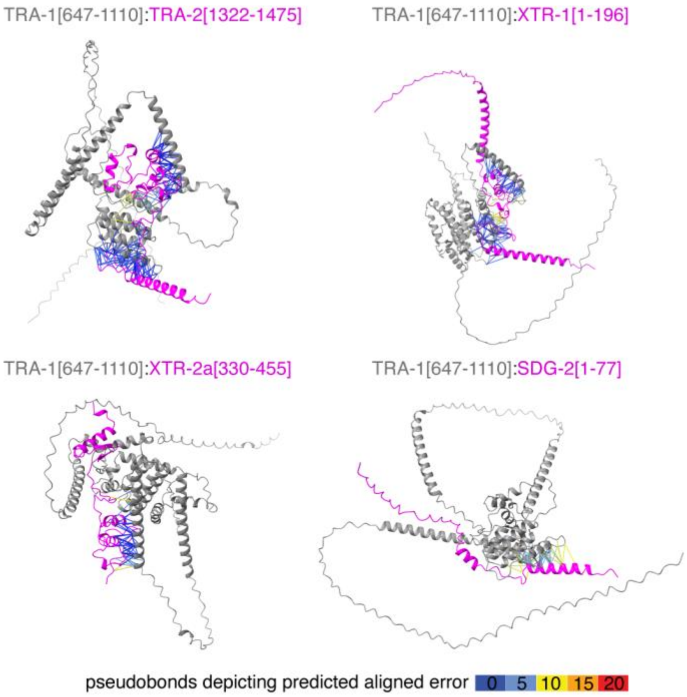


**Suppl. Fig. 8.** Related to Fig. 5. Predicted interactions of TRA-1 with TRA-2 and regions from related proteins, including SDG-2. Numbers indicate amino acid residues included for the structure prediction using AlphaFold3. Colors and pseudobonds are as in Fig. 5c.

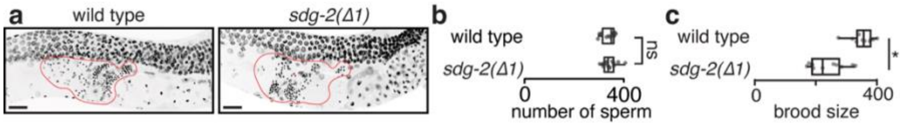


**Suppl. Fig. 9.** Related to Fig. 6. Different cohorts of *sdg-2(Δ1)* animals could have different numbers of progeny. a) Representative images showing spermatheca region (red outline) with sperm (small punctae) in wild-type (left) and *sdg-2(Δ1)* animals. Scale bar, 20 µm. b) Box plots showing numbers of sperm measured after DAPI staining of wild-type animals and one cohort of *sdg-2(Δ1)* animals, consistent with the normal brood size as measured in Fig. 6e, top. c) Box plots showing the brood size of a different generation of *sdg-2(Δ1)* animals measured along with paired wild-type animals assayed at the same time, replotted from Fig. 6e, bottom. Asterisks and n are as in Fig. 4b.

**Supplementary Files**

**File 1.** List of genes changed in *sid-1* mutants as detected using RNA-seq. [excel]

**File 2.** Background mutations detected using whole-genome sequencing in animals subject to genome editing of *sid-1*. [excel]

**File 3.** List of proteins with structural similarity to the predicted structure of SDG-1 as detected by VAST. [excel]

**File 4.** Localization of SDG-1::mCherry to germ granules and/or microtubules under different conditions. [excel]
